# A DNA-origami nuclear pore mimic reveals nuclear entry mechanisms of HIV-1 capsid

**DOI:** 10.1101/2020.08.10.245522

**Authors:** Qi Shen, Chaoyi Xu, Sooin Jang, Qiancheng Xiong, Swapnil C. Devarkar, Taoran Tian, Gregory J. Bedwell, Therese N. Tripler, Yingxia Hu, Shuai Yuan, Joshua Temple, Jiong Shi, Christopher Aiken, Alan N. Engelman, Juan R. Perilla, C. Patrick Lusk, Chenxiang Lin, Yong Xiong

**Affiliations:** Department of Molecular Biophysics and Biochemistry, Yale University, New Haven, CT, 06511, USA; Department of Cell Biology, Yale School of Medicine, New Haven, CT, 06520, USA; Nanobiology Institute, Yale University, West Haven, CT, 06516, USA; Department of Chemistry and Biochemistry, University of Delaware, Newark, DE, 19716, USA; Department of Cancer Immunology and Virology, Dana-Farber Cancer Institute, Boston, MA, 02215, USA; Department of Medicine, Harvard Medical School, Boston, MA, 02115, USA; Department of Pathology, Microbiology and Immunology, Vanderbilt University Medical Center, Nashville, TN, 37232, USA

**Keywords:** HIV-1 capsid, nuclear entry, DNA-origami, nuclear pore mimic, RRR-motif, pattern sensing, curvature preference, capsid stabilization, capsid deformation

## Abstract

The capsid of human immunodeficiency virus 1 (HIV-1) plays a pivotal role in viral nuclear import, but the mechanism by which the viral core passages the nuclear pore complex (NPC) is poorly understood. Here, we use DNA-origami mimics of the NPC, termed NuPODs (NucleoPorins Organized by DNA), to reveal the mechanistic underpinnings of HIV-1 capsid nuclear entry. We found that trimeric interface formed via three capsid protein hexamers is targeted by a triple-arginine (RRR) motif but not the canonical phenylalanine-glycine (FG) motif of NUP153. As NUP153 is located on the nuclear face of the NPC, this result implies that the assembled capsid must cross the NPC *in vivo*. This hypothesis is corroborated by our observations of tubular capsid assemblies penetrating through NUP153 NuPODs. NUP153 prefers to bind highly curved capsid assemblies including those found at the tips of viral cores, thereby facilitating capsid insertion into the NPC. Furthermore, a balance of capsid stabilization by NUP153 and deformation by CPSF6, along with other cellular factors, may allow for the intact capsid to pass NPCs of various sizes. The NuPOD system serves as a unique tool for unraveling the previously elusive mechanisms of nuclear import of HIV-1 and other viruses.

## Introduction

As a retrovirus, HIV-1 reverse-transcribes its RNA genome for integration into host chromatin within the nucleus (Lusic and Siliciano, 2017). However, it remains unclear how HIV-1 gains access to the nuclear compartment, which is segregated from the cytoplasm by a double membraned nuclear envelope. The most logical hypothesis is that HIV-1 passes through nuclear pore complexes (NPCs), which are massive protein assemblies embedded in the nuclear envelope that control molecular exchange between the nucleus and the cytoplasm. Indeed, recent evidence suggests that the intact HIV-1 capsid is capable of crossing the NPC and disassembles only in the nuclear interior (Burdick et al., 2020). However, this is conceptually challenging, given that the commonly accepted ∼40 nm diameter of the NPC central transport channel (von Appen et al., 2015) is smaller than the ∼60 nm width of an intact capsid (Briggs et al., 2003; Mattei et al., 2016). While recent data supports that there may be energy-dependent changes in the NPC scaffold that might dilate the channel (Zimmerli et al., 2020), the channel itself is also filled with dozens of intrinsically disordered proteins (nucleoporins/NUPs) rich in repeating FG motifs, which establish a diffusion barrier for macromolecules. The mechanism by which a largely intact HIV-1 capsid passes through the NPC central channel remains to be elucidated.

HIV-1 capsid is composed of capsomeres of approximately 250 capsid protein (CA) hexamers and 12 CA pentamers, which assemble into a fullerene cone (Briggs et al., 2003; Ganser et al., 1999; Li et al., 2000; Mattei et al., 2016; Pornillos et al., 2011). The capsid furnishes an enclosed compartment necessary for reverse transcription and concomitantly protects the genome from destruction by host immune surveillance and cellular antiviral restriction factors (Hulme et al., 2015; Hulme et al., 2011; Yamashita and Engelman, 2017). Most capsid-targeting host factors sense higher-order patterns formed by multiple capsomeres, and *in vitro* assembly techniques have matured in recent years to capture such patterns. CA can be assembled through various disulfide bonds into standalone hexamers or pentamers (Pornillos et al., 2009; Pornillos et al., 2011), or multi-hexamer/pentamer units as well as a hexamer-2 structure that mimics the interface of three hexamers in the assembled capsid (Summers et al., 2019). Further assembly of higher-order structures such as CA tubes (Lopez et al., 2011), spheres (Zhang et al., 2018), and stable cones are also achievable (Dick et al., 2018) (Figure 1A). However, these assemblies have not yet been applied to exploring interactions with the NPC as there are few *in vitro* platforms that can mimic the dimensions and complexity of the NPC.

**Figure 1.**
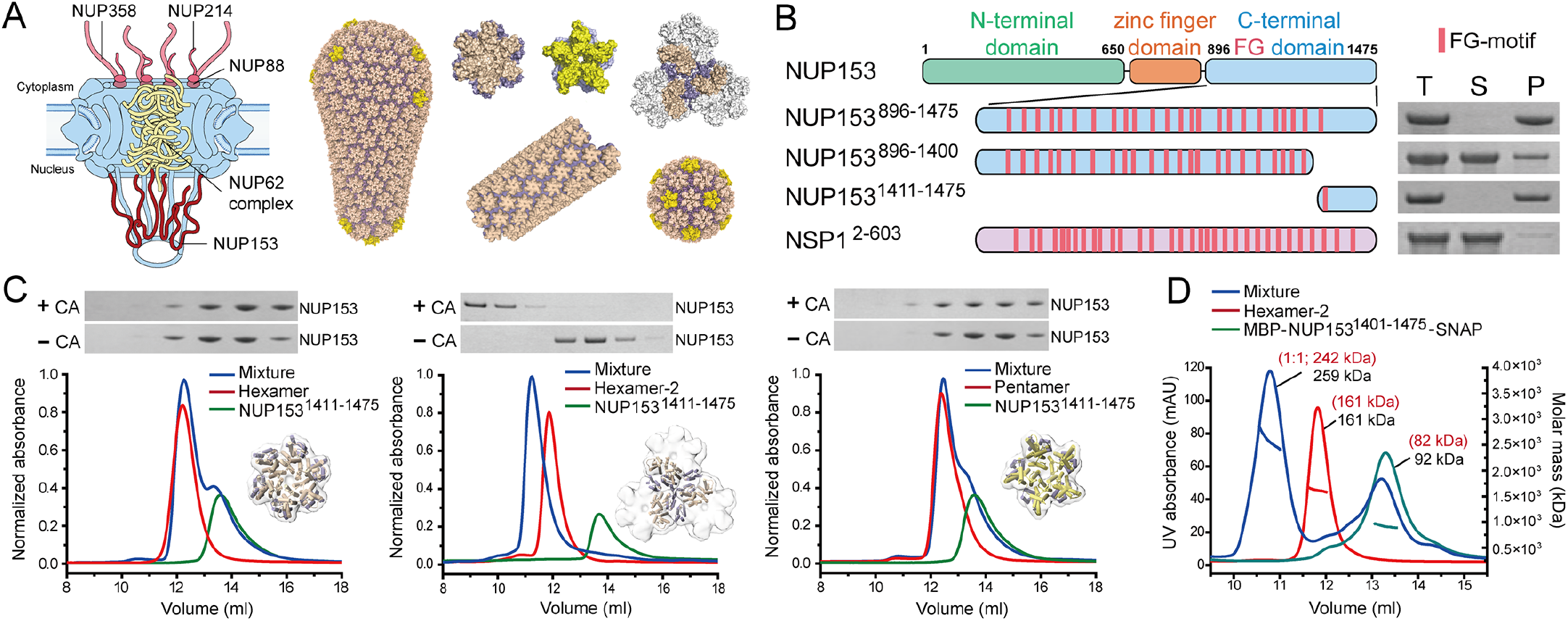
NUP153 targets the lattice interfaces in the assembled capsid. (A) Schematic of the NPC with NUPs that have been shown to bind to HIV-1 capsid (left) and various *in vitro* assembled CA structures used in this study (right). (B) Co-pelleting assay of NUP153 CTD variants and yeast nucleoporin NSP1 control with CA tubes. Left: Schematics of the constructs with FG repeats in each protein indicated as red bars. Right: The Total (T), Soluble (S), and Pellet (P) fractions of co-pelleting assays using A14C/E45C disulfide crosslinked CA tubes analyzed by SDS-PAGE. (C) Size-exclusion chromatography co-elution assays of CA hexamer (left), hexamer-2 (middle) or pentamer (right) with NUP153^1411-1475^. Top: SDS-PAGE of eluates. Only hexamer-2 induced a peak shift of NUP153^1411-1475^, indicating co-elution, with a corresponding shift of the elution fraction bands in SDS-PAGE (top). (D) SEC-MALS analysis of NUP153^1401-1475^ and hexamer-2. Overlay of the individual SEC-MALS elution profiles of NUP153^1401-1475^ (red), hexamer-2 (green), and a mixture of the two (blue). The measured (black) and predicted (red) molecular weights are shown. See also Figure S1.

At ∼100 MDa, the NPC is the largest macromolecular complex in the human cell, which is built from ∼30 different NUPs each in 16 or 32 copies (Brohawn et al., 2009; Cook et al., 2007; Lin and Hoelz, 2019a; Strambio-De-Castillia et al., 2010). Many FG-rich NUPs can interact with HIV-1 capsid, including NUP358, NUP214, NUP88, NUP62, and NUP153 (Bichel et al., 2013; Dharan et al., 2020; Di Nunzio et al., 2013; Kane et al., 2018; Matreyek and Engelman, 2011). Apart from these NUPs, soluble nuclear transport receptors (NTRs) including Importin α3 (Ao et al., 2010), Karyopherin β2/Transportin-1 (TRN-1) (Fernandez et al., 2019), and Transportin-3 (TRN-SR2) (Maertens et al., 2014; Valle-Casuso et al., 2012) are implicated in HIV-1 nuclear entry. Other host factors, including cyclophilin A (CypA) (Schaller et al., 2011; Yamashita and Engelman, 2017), cleavage and polyadenylation specificity factor 6 (CPSF6) (Achuthan et al., 2018; Bejarano et al., 2019; Bhattacharya et al., 2014; Chin et al., 2015; Price et al., 2012; Sowd et al., 2016), and myxovirus resistance protein B (MxB) (Busnadiego et al., 2014; Kane et al., 2018; Kane et al., 2013; King et al., 2004), can bind the capsid and affect its nuclear transport. This complexity limits the impact of cellular methods such as targeted NUP knockdowns to reveal capsid transport mechanisms (Kane et al., 2018).

To establish a simplified model system that circumvents these difficulties, we took advantage of our prior generation of the NucleoPorins Organized by DNA (NuPOD) platform (Fisher et al., 2018). This experimental system leverages the DNA-origami technique (Dietz et al., 2009; Rothemund, 2006; Seeman and Sleiman, 2018) to assemble a DNA ring that mimics the ∼40 nm diameter of the NPC transport channel (Fisher et al., 2018; Ketterer et al., 2018; von Appen et al., 2015). Multiple copies of multiple NUPs can be attached to the DNA-origami rings at specific positions, with additional tunable parameters such as protein stoichiometry, grafting density, and topology (Fisher et al., 2018; Ketterer et al., 2018). Therefore, our NuPOD system is ideally suited to interrogate the involvement of individual or specific combinations of NUPs in HIV-1 capsid nuclear import. Here we have focused on NUP153, a component of the NPC’s nuclear basket (Schmitz et al., 2010; Walther et al., 2001), because it binds capsid directly and is essential for HIV-1 capsid nuclear transport (Matreyek and Engelman, 2011; Price et al., 2014). Since NUP153 resides in the nuclear side of the NPC (Buffone et al., 2018; Duheron et al., 2014), the capsid would need to insert deep into or pass through the NPC to reach it. Furthermore, as NUP153 plays an important role in the late stage of capsid nuclear transport, understanding its actions is key to unraveling the nuclear entry mechanisms of HIV-1 (Chen et al., 2016; Matreyek and Engelman, 2011; Matreyek et al., 2013).

In this work, we combined NuPOD with capsid engineering techniques to reveal the mechanistic underpinnings of capsid nuclear import *in vitro*. We identified the mode of capsid pattern-sensing by the NUP153 C-terminal tail, via an RRR-motif but not the canonical FG-motif, which strongly binds higher-order capsid structures beyond single hexamers and in doing so stabilizes the CA lattice. We created NuPODs with 32 copies of the NUP153 C-terminal domain (residues 896-1475; NUP153CTD), the endogenous number in the NPC (Ori et al., 2013), to simulate the critical features of the natural nuclear pore environment. Capsids isolated from HIV-1 virions and *in vitro* assembled structures bind these NUP153 NuPODs in a shape and curvature dependent manner. Strikingly, tubular CA assemblies can thread through the central pore of the NUP153 NuPOD, strongly supporting the transport of the assembled viral capsid through the nuclear pore.

## Results

### NUP153 targets lattice interfaces of the assembled capsid

It is well established that NUP153 directly interacts with the capsid and is essential for HIV-1 nuclear import. NUP153 is composed of 1475 amino acids (aa) and contains 24 FG repeats in its CTD (Figure 1B) (Fahrenkrog et al., 2002; Matreyek et al., 2013). While the CA hexamer contains sites for general FG-motif binding, it has been shown that only the last FG-motif in the C-terminal tail of NUP153 was needed for the capsid interaction (Matreyek et al., 2013). Consistently, crystal structures showed that the FG-motif in the NUP153 C-terminal tail binds at the interface between two subunits within the CA hexamer, albeit with a relatively weak affinity (*K*_*d*_ of ∼50 μM) (Buffone et al., 2018; Price et al., 2014).

We tested the interaction between purified NUP153 constructs and nanotubes assembled from purified CA protein in a co-pelleting assay (Figure 1B and S1). For this purpose, A14C/E45C disulfide cross-linked CA tubes were employed due to their known stability under low salt conditions (Lopez et al., 2011). Consistent with previous findings, the C-terminal tail (residues 1411–1475, NUP153^1411-1475^) efficiently bound to CA tubes. The preceding NUP153 region (residues 896-1410 with 23 FG-repeats) or control NUP, yeast nucleoporin NSP1 (*S. cerevisiae* orthologue of NUP62 with 32 FG-repeats) showed little or no binding, respectively. These results, together with the reported low binding affinity of the NUP153 FG-containing peptide to CA hexamers (*K*_*d*_ ∼50 μM), highlights that FG motifs have limited inherent capsid-binding activity. By contrast, we observed complete co-pelleting of NUP153^1411-1475^ with CA tubes even at comparatively low NUP concentrations (3 μM) (Figure 1B and S1). Since the CA tubes present the higher-order lattice of the assembled capsid, it suggests the existence of a strong interaction beyond the hexamer-specific NUP153 FG-motif binding site.

To provide a more detailed understanding of how NUP153 binds HIV-1 capsid, we analyzed the interaction between NUP153^1411-1475^ and various CA assemblies representing different patterns on the capsid. We performed size exclusion chromatography (SEC) analysis of NUP153^1411-1475^ with individual capsomeres (CA hexamers or pentamers), as well as the hexamer-2 assembly that mimics the interface between 3 hexamers (Summers et al., 2019) (Figure 1C). Consistent with the reported low binding affinities (Price et al., 2014), neither CA hexamer nor pentamer detectably bound NUP153^1411-1475^. By contrast, efficient binding of hexamer-2 to NUP153^1411-1475^ was readily detected (Figure 1C). We further used SEC-MALS (SEC-multiangle light scattering) to measure the molecular weight distributions in each elution peak and deduce the stoichiometric ratios of NUP153 and hexamer-2 in the complex. The results showed good agreement with a 1:1 molar ratio of NUP153^1401-1475^: hexamer-2 (Figure 1D), pointing to a model of the C-terminal tail of NUP153 engaging inter-hexamer interfaces on the capsid. These observations revealed that NUP153 preferentially engages the higher-order lattice within the assembled capsid, implying that capsid interaction with the NPC requires at least a partially assembled capsid.

### The positively charged C-terminal tail of NUP153 binds to the 3-fold tri-hexamer interface in capsid

The higher-order capsid lattice preference of NUP153^1411-1475^ suggested it contains a major binding site beyond the CA hexamer-targeting FG-motif. We performed a comprehensive mapping study to identify the specific region responsible for high-affinity binding with the capsid (Figure 2A and S2). Substantial binding (60%-75%) remained with deletion or truncation of the FG-motif (NUP153^1401-1475FTFGΔ^ or NUP153^1425-1475^), supporting the notion that the FG-motif provides some but not the majority of the binding affinity (Figure 2A and S2). When the N-terminal region of the NUP153 C-terminal tail was progressively removed (NUP153^1436-1475^), a moderate further decrease in CA tube binding was observed. Interestingly, even the shortest C-terminal tail construct, containing only the last 11 aa of the protein (NUP153^1465-1475^), retained close to 50% binding. On the other hand, removal of the last 11aa region (NUP153^1411-1465^) retained only ∼25% of the binding. These results indicated that the C-terminal 11 residues of NUP153 contain a strong capsid-binding site (Figure 2A and S2).

**Figure 2.**
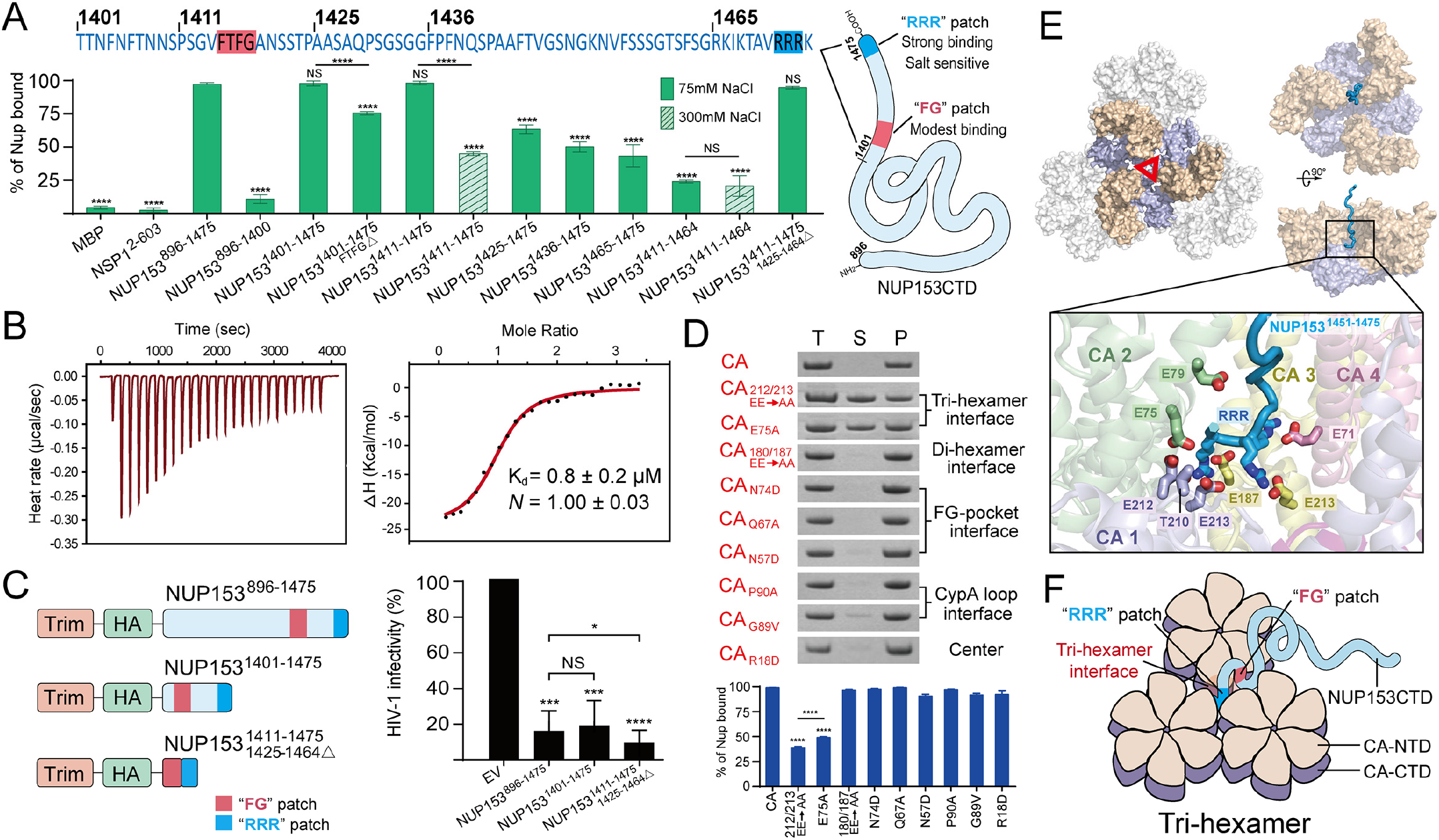
A bipartite interaction between NUP153 and CA lattice. (A) Top: The C-terminal 75 aa of NUP153. Bottom: percentages of various NUP153 variants, along with maltose-binding protein (MBP) and yeast nucleoporin NSP1 negative controls, bound to CA tubes in the co-pelleting assays under either low salt (75 mM NaCl, solid green bar) or high salt (300 mM NaCl, striped green bar) conditions. Data were expressed as mean ± standard deviation. Differences were tested by two-tailed Student’s t-test. Compared with NUP153^894-1475^; NS, not significant (*P*⩾0.05); ****, *P*<0.0001. The schematic of the two capsid-binding motifs on NUP153CTD is shown in the right panel. (B) The NUP153^1411-1475/1425-1464Δ^ peptide (25 aa) binds to CA hexamer-2 with 0.8 ± 0.2 μM *K*_*d*_ as measured by ITC. (C) Restriction of HIV-1 infection by TRIM-NUP153 fusion proteins. Schematic of test constructs (left panel) and results (avg. ± SD of two independent experiments) of HIV-1 infection assays (right panel) are shown. EV, empty vector. Statistical significance was determined by one-tailed Student’s t-test; NS, not significant (*P*≥0.05); *, *P*<0.05; ***, *P*<0.001; ****, *P*<0.0001. Immunoblot of protein expression is shown in Figure S2C. (D) Co-pelleting assays of CA mutants with NUP153^1411-1475^. Top: SDS-PAGE analysis of co-pelleted NUP153^1411-1475^. Bottom: quantification of bound NUP153^1411-1475^ normalized to that of wild-type CA tubes. Data were expressed as mean ± standard deviation. Differences were tested by two-tailed Student’s t-test. Compared with A14C/E45C CA tubes; ****, *P*<0.0001. (E) All-atom molecular dynamic simulations identify NUP153CTD interactions at the CA tri-hexamer center. Top: surface representation of the hexamer-2 tri-hexamer center region (left) and simulated NUP153^1451-1475^ (light blue cartoon) binding to hexamer-2 (right). Bottom: the RRR-motif is surrounded by negatively charged glutamate residues (sticks) from 4 CA monomers (bottom). (F) Schematics of the bipartite interaction between NUP153CTD and the CA tri-hexamer. See also Figure S2, Figure S3, Table S1, and Movie S1.

The short 11 aa C-terminal tail of NUP153 is enriched in positively-charged amino acids, including a triple-arginine (RRR) motif that bears resemblance to the capsid-binding motif in MxB (Busnadiego et al., 2014; Kane et al., 2018; Kane et al., 2013; King et al., 2004). Since the MxB RRR-motif targets capsid through electrostatic interactions, we tested the effect of increasing salt concentrations on CA tube binding. Indeed, when the NaCl concentration was increased from 75 mM to 300 mM, NUP153^1411-1475^ lost over half of its binding capacity, while NUP153^1411-1464^ containing the FG-motif but not the 11 aa tail showed no change in its binding (Figure 2A and S2). This salt sensitivity suggests that the RRR-motif engages the capsid in an MxB-like manner, likely binding at the negatively charged tri-hexamer center.

To test the hypothesis that NUP153 binds the capsid in a bipartite mode using both the FG-motif and the RRR-motif, we created a construct connecting the FG-motif to the last 11 aa region (NUP153^1411-1475/1425-1464Δ^). The resultant 25 aa segment bound CA nanotubes as efficiently as the NUP153CTD (NUP153^896-1475^) did, confirming these two regions provide most of the capsid-binding capacity. We further used isothermal titration calorimetry (ITC) to quantify the binding affinity of this 25 aa peptide to the lattice-mimicking hexamer-2. This minimal NUP153 CTD peptide bound to hexamer-2 with a *K*_d_ of ∼0.8 μM, which is close to two orders of magnitude tighter than the *K*_*d*_ of ∼50 μM reported for FG-hexamer binding (Price et al., 2014) (Figure 2B).

To confirm the results of *in vitro* binding assays under HIV-1 infection conditions, we employed a previously-described tripartite motif (TRIM) construct wherein the C-terminal domain of NUP153^896-1475^ was fused to the RING, B-box 2, and coiled coil domains of rhesus TRIM5α (Matreyek et al., 2013). Effector functions provided by the TRIM domains coupled with capsid binding, which was provided by the NUP153CTD, significantly inhibited HIV-1 infection (Matreyek et al., 2013) (Figure 2C). Deletion mutagenesis revealed that TRIM-NUP153^1401-1475^ lacking the 505 N-terminal residues of 896-1475, as well as a minimal fusion construct that harbored just the 25 residues composed of the FG-motif (1411-1424) and RRR-motif (1465-1475), conferred full restriction activity (Figure 2C). These results support the findings from biochemical studies and confirm that NUP153 associates with the capsid via two regions: the first is the previously reported weaker FG-hexamer interface; the second is the positively charged RRR-motif that binds strongly (Figure 2A, F).

To identify the capsid surface that is targeted by the RRR-motif of NUP153, we mutated key CA residues that are located in different regions of the capsid, including the tri-hexamer interface (EE212/213AA, E75A), the di-hexamer interface (EE180/181AA), the FG pocket (N74D, Q67A, N57D), the CypA-binding loop (P90A, G89V), and the hexamer central pore (R18D) (Bhattacharya et al., 2014; Dick et al., 2018; Price et al., 2014; Schaller et al., 2011; Smaga et al., 2019). Co-pelleting results showed that alterations of key residues at the tri-hexamer interface significantly reduced NUP153 binding to CA nanotubes (Figure 2D). By contrast, mutations in the di-hexamer interface, the CypA-binding loop, the FG-binding pocket, and the hexamer central pore showed little or no effect on binding (Figure 2D). These data again demonstrate the strong affinity of NUP153 for binding the CA lattice beyond just the hexamer. Similar to MxB’s interaction with the CA lattice, the main driving force of association is the electrostatic contact between the positively charged RRR motif and the negatively charged CA tri-hexamer interface (Smaga et al., 2019).

We next performed all-atom molecular dynamics (MD) simulations to gain a mechanistic understanding of the NUP153 CTD-CA interactions at the tri-hexamer interface. A NUP153^1451-1475^ and hexamer-2 complex model was built and simulated for 8.7 μs. The results of this MD simulation are shown in Figure S3 and Table S1. NUP153^1451-1475^ was found to be mostly unstructured during the simulations (Figure S3A), except for a short α-helix at the C-terminus (residue 1470 to 1475) that appeared temporarily (Figure S3C and Movie S1). The interaction between NUP153^1451-1475^ and hexamer-2 appeared to be dynamic, rapidly transitioning between bound and unbound states as analyzed by the Markov state model (Chodera and Noe, 2014; Husic and Pande, 2018a; Wang et al., 2018b) (Figure S3). The NUP153 C-terminal tail formed high occupancy contacts with polar CA residues in different states (Table S1) whereby in the last state the RRR motif inserted deeply into the tri-hexamer interface, surrounded by negatively charged glutamate residues from 4 different CA monomers (Figure 2E, bottom).

### Building a DNA-origami mimic of NUP153 organization within the human NPC

To interrogate capsid-NUP153 interaction in a scenario resembling the natural environment, we generated NuPODs with NUP153CTD attached to the inner surface of the 45 nm diameter DNA-origami ring (Fisher et al., 2018). The inner side of the ring has 32 single-stranded DNA “handles”, to which complementary DNA oligonucleotides (anti-handles) conjugated to NUP153CTD can hybridize. We constructed 32 handles inside the NuPOD ring since there are 32 copies of NUP153 in the human NPC (Lin and Hoelz, 2019b; Ori et al., 2013). NUP153CTD was fused to a SNAP-tag, which reacts with a benzylguanine-labeled DNA oligo (BG-oligo) to covalently link the anti-handle to the protein. The SNAP-tag/BG-oligo reaction was very efficient, yielding more than 95% of NUP153CTD conjugation with anti-handles (Figure 3A). NUP153CTD-conjugated anti-handles were then hybridized with complementary handles at the inside of the origami ring to produce the NUP153 NuPOD, which was then purified by rate-zonal centrifugation (Lin et al., 2013). We used negative-stain electron microscopy (EM) to confirm the NUP153 NuPOD construction, which was characterized by NUP153CTD filling the origami ring interior (Figure 3B and S4).

**Figure 3.**
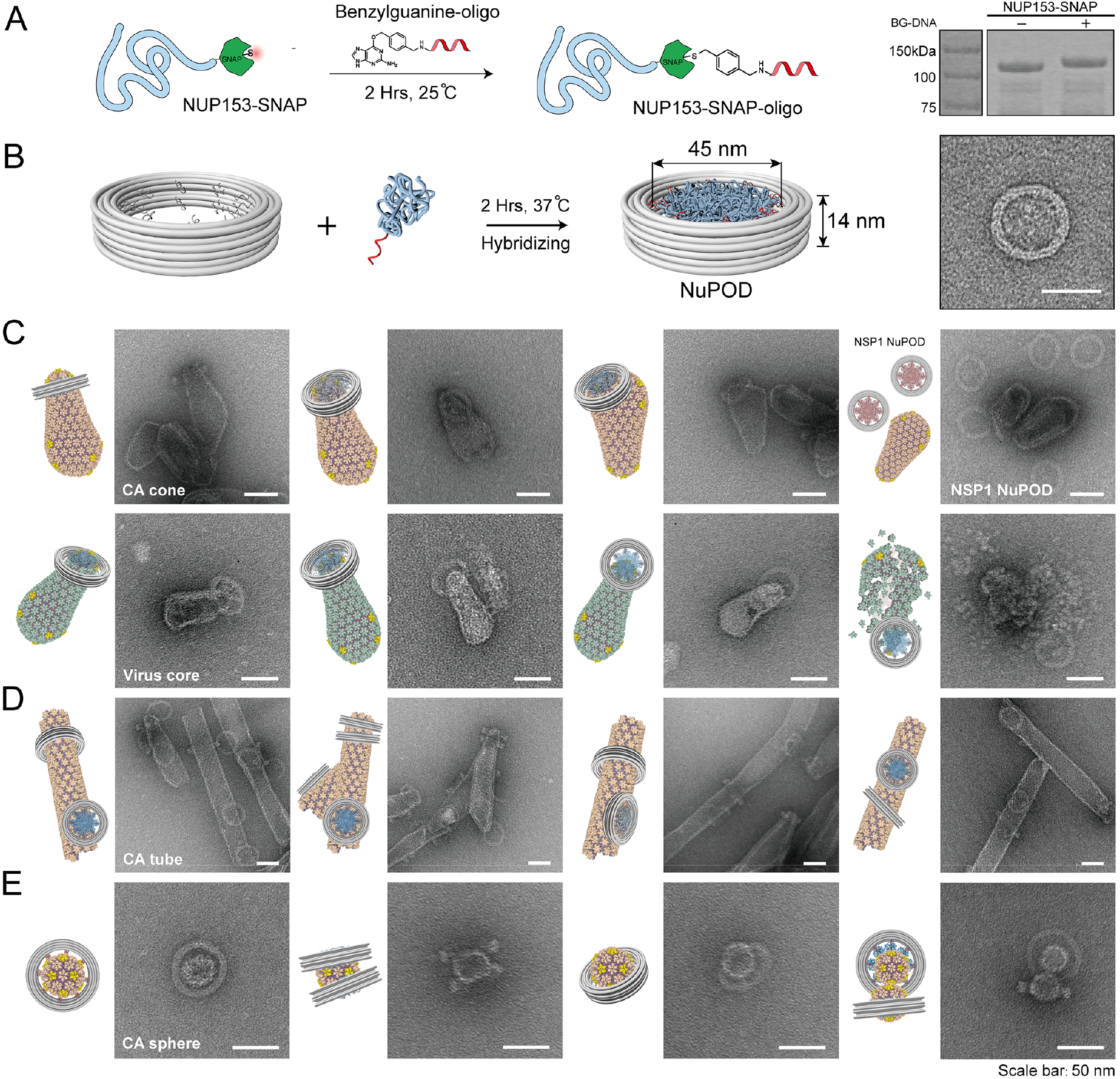
NUP153 NuPOD binding to capsid assemblies. (A) Schematic of conjugating anti-handle (BG-DNA oligonucleotide) to NUP153-SNAP (left). Right: SDS-PAGE analysis of the conjugation product compared to the original unconjugated protein. Note the shift in band positions. (B) Construction of NuPOD to mimic critical features of the NPC. Left: schematic of NUP153-SNAP-oligo hybridization to a DNA origami ring. Right: negative-stain electron micrograph of a NUP153CTD grafted NuPOD. Scale bars: 50 nm. (C) Schematics and negative-stain electron micrographs of the NUP153 NuPOD bound to *in vitro* assembled capsid cones (top) and purified native virus cores (bottom). Assembled capsid cones did not detectably bind to the yeast nucleoporin NSP1 NuPOD (top right). The NUP153 NuPOD also bound partially disassembled native cores (bottom right). Scale bars: 50 nm. (D) Schematics and negative-stain electron micrographs of the NUP153 NuPOD bound to CA tubes, either on side surfaces or with CA tubes threading through the NuPOD. Scale bars: 50 nm. (E) Schematics and negative-stain electron micrographs of the NUP153 NuPOD bound to CA spheres. Scale bars: 50 nm. See also Figure S4.

### NUP153 NuPOD binds to the tip regions of *in vitro* assembled capsids and native viral cores

We utilized the NUP153 NuPOD to gain mechanistic insights into the NPC engagement of the cone-shaped HIV-1 capsid. We first tested its interaction with *in vitro* assembled CA cones (average length ∼120 nm, width ∼60 nm), which were produced using the A14C/E45C CA mutant in the presence of inositol hexakisphosphate (IP6) and CypA (Dick et al., 2018). As expected, the A14C/E45C crosslinked cones remained stable under low salt conditions (Figure S4). As visualized by negative-stain EM, the cones readily bound the NUP153 NuPODs. Under these conditions, the broad or narrow curved tip regions of the capsid cones were found to exclusively insert into the NUP153 NuPODs. We did not observe any NuPOD attachment to the flat surface of the capsid cone. NuPODs assembled with yeast nucleoporin NSP1 served as a negative control. Consistent with its lack of co-pelleting with CA nanotubes (Figure 2A), capsid cones failed to bind the yeast nucleoporin NSP1 NuPODs (Figure 3C, top right).

We next tested the binding of native virus cores with the NUP153 NuPOD. Native cores were prepared from concentrated HIV-1 particles by sucrose density gradient centrifugation (Shah and Aiken, 2011). The results were similar to those obtained using *in vitro* assembled capsid cones; the tip regions of the native cores could bind and insert into the NUP153 NuPOD (Figure 3C). These findings suggest that NUP153 may have a curvature preference, specifically targeting the tips of HIV-1 capsid (described more below). Interestingly, the native core preparations contained partially dissolved cores, which likely contain deformed or partially disassembled capsids. We also observed NuPOD rings binding these partially disassembled structures. Because NUP153 targets higher-order tri-hexameric interfaces on the capsid (Figure 1 and 2), these partially dissolved cores likely retained this aspect of the lattice structure (Figure 3C, bottom right).

### CA tubes bind and thread through the NUP153 NuPOD

We also investigated the interaction of the NUP153 NuPOD with CA tubes. As above, A14C/E45C cross-linked CA tubes (diameter 40-50 nm) were used due to their stability under low salt conditions (Lopez et al., 2011) (Figure S4). NUP153 NuPOD readily bound CA tubes, often on the flat side surfaces (Figure 3D). Strikingly, we also observed CA tubes completely penetrating the entire depth of and threading through the NuPOD rings, sometimes with more than one attached at the tube tip (Figure 3D). This result provides direct evidence of the ability of intact capsid assemblies to pass through the central channel of a nuclear pore-like confinement. The passage of CA tubes presumably occurred for those with a diameter less than that of the NuPOD, although our data cannot tell if there was local constriction caused by NUP153 at the site of contact. We observed that the NUP153 NuPOD rings “glide” far along CA tubes, indicating that NUP153 can diffuse along the CA lattice. It is not clear whether there is directionality to this diffusion.

### NUP153 prefers to bind regions of high capsid curvature

The observed dominant tip-binding mode of capsid cones to NUP153 NuPODs prompted us to investigate whether NUP153 has a curvature preference for binding. The cone-shaped capsid has a “flat” surface on its sides and highly curved tip regions (Liu et al., 2015) (Figure 4A). It is well established that the CA tube is a good mimic of the flat side of the capsid (Zhao et al., 2011). Additionally, it was reported that the N21C/A22C CA mutant can assemble into CA spheres (30-45 nm) that mimic the highly curved capsid tips (Zhang et al., 2018; Zhao et al., 2011) (Figure 4A and S4). We assembled N21C/A22C CA spheres and confirmed that these bound well to the NUP153 NuPOD, mostly residing in the middle of the NuPOD rings. When the NuPODs were in excess, we observed two NuPOD rings attaching to the same CA sphere (Figure 3E).

**Figure 4.**
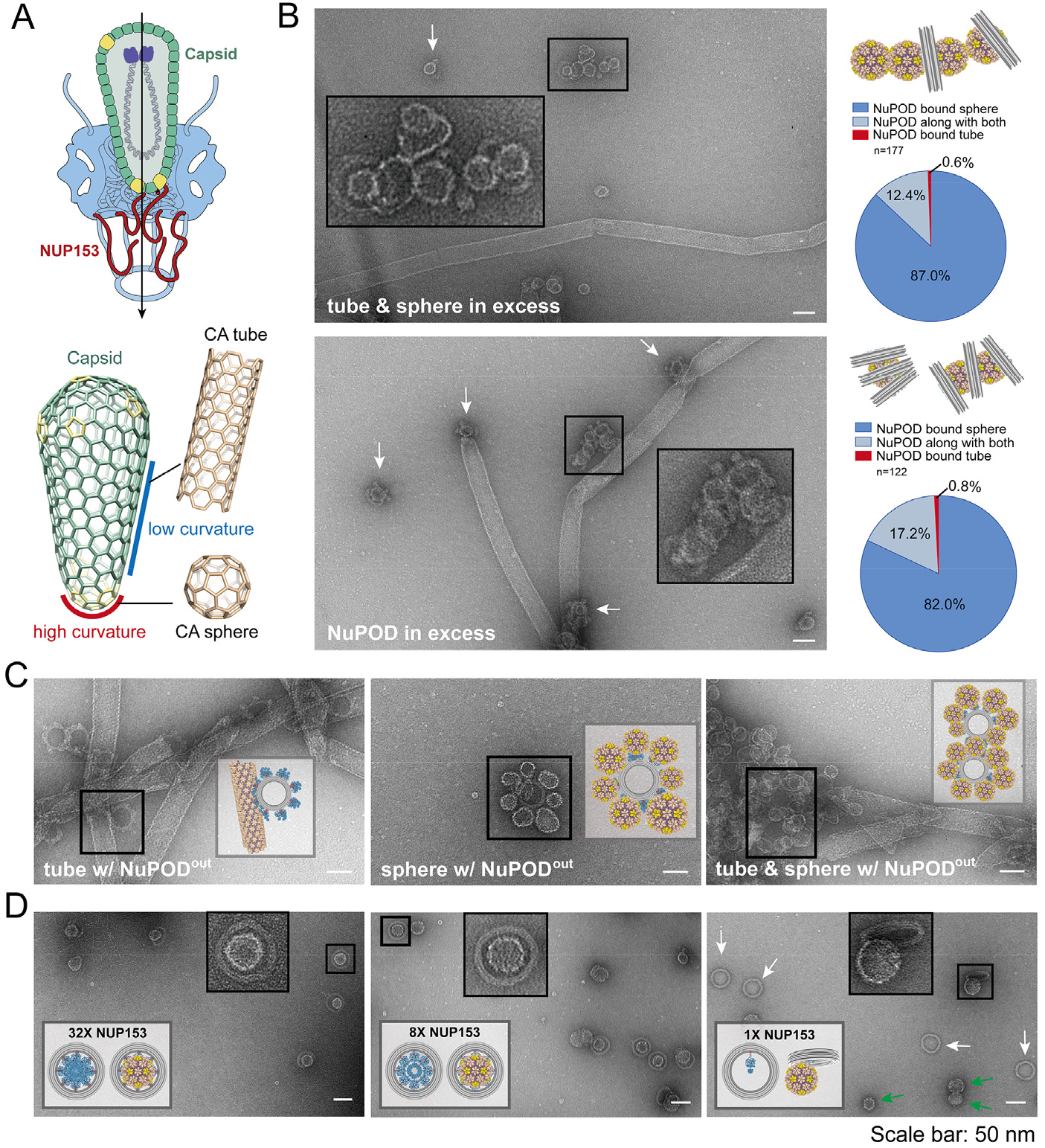
NUP153 prefers highly-curved regions of capsid for avid NuPOD association. (A) Top: Schematic of capsid tip insertion into the NPC central pore during HIV-1 nuclear transport. Bottom: Schematics showing that the capsid cone has a flat surface on its sides and higher curvature at the tips. CA tubes mimic the sides of the capsid and CA spheres resemble the tips. (B) Left: Negative-stain electron micrographs of NUP153 NuPOD binding to a 1:1 mixture of CA spheres and tubes in excess (top) versus binding when the NuPOD was in excess (bottom). The white arrows and insets mark the sphere-bound NuPODs. Right: Schematics of the major binding mode and quantification of all binding events. Quantifications suggest a sphere-preferred binding of NuPOD (Statistical significances were determined by the Chi-Squared Test, P<0.0001). Scale bars: 50 nm. (C) Negative-stain electron micrographs of the external-grafted NUP153^out^ NuPOD binding to CA tubes alone (left), CA spheres alone (middle), and a 1:1 mixture of CA spheres and tubes (right). Note the lack of CA sphere binding to the flat tube surface in the mixture experiment. Scale bars: 50 nm. (D) Negative-stain electron micrographs of CA spheres binding to NuPODs with 32 (left), 8 (middle), or 1 (right) copies of NUP153CTD. Most of the single-NUP153 NuPOD remain empty without bound CA spheres (right, marked with white arrows). The free spheres are marked with green arrows. Scale bars: 50 nm.

CA tube and sphere assemblies capture two different surface geometries present in native capsids. We accordingly used these assemblies in binding competition assays to elucidate the curvature preference of NUP153. NUP153 NuPODs challenged with equimolar mixtures of CA spheres and tubes were analyzed by negative-stain EM (Figure 4B). We first applied a tube:sphere mixture (1:1 ratio) in excess of the NuPODs, and observed that almost all NuPODs bound to CA spheres, including some NuPODs with two spheres attached. Quantification revealed that close to 90% of the NuPODs bound CA spheres only, compared to ∼0.5% bound to CA tubes, which was over a 100-fold difference (Figure 4B, bottom left). When NuPODs were added in excess of the CA sphere:tube mixture (1:1 ratio), we again found over a 100-fold preference for sphere binding over tubes (Figure 4B, bottom right). The excess NuPODs did not lead to increased binding to CA tubes, but rather greater NuPOD occupancy (up to three) bound to the same CA sphere. As the CA spheres mimic the capsid tip geometry, this is consistent with our observation that the NUP153 NuPOD preferentially bound to the tips of either *in vitro* assembled capsid cones or purified virus cores (Figure 3C).

We note that because NUP153 was positioned inside the NuPOD ring, the observed CA sphere preference can be potentially due to geometric constraints leading to greater inaccessibility of CA tubes. To exclude this possibility, we repositioned the DNA handles to the exterior of the origami ring to create a NuPOD with fully exposed NUP153CTD. This exterior-grafted NUP153 NuPOD (NUP153^out^ NuPOD) efficiently bound CA tubes (on the flat side) and spheres (6-8 spheres per NuPOD) in tube-only or sphere-only reactions (Figure 4C). Consistent with the earlier competition results, the majority of NUP153^out^ NuPODs clustered around the CA spheres in the mixture experiment, without binding to the flat surface of CA tubes (Figure 4C). These results confirm that the observed preference for capsid-tip curvature is an intrinsic property of NUP153. It is conceivable that this preference facilitates the directional transport of the capsid through the NPC.

### Multiple copies of NUP153 provide binding avidity to the capsid

As there are 32 copies of NUP153 in a natural NPC, we constructed NuPODs with varying numbers of NUP153CTD attached (1, 8, or 32) to understand how binding avidity affects the association of the capsid to the NPC (Figure 4D). NuPODs with either 8 or 32 NUP153 copies adhered robustly to CA spheres. By contrast, NuPODs with only 1 copy of NUP153 bound CA spheres poorly, with only ∼10% of NuPODs having attached spheres (Figure 4D). These results suggest that a single NUP153 molecule is insufficient for stable capsid attachment to the NPC and that binding avidity from multiple copies of NUP153 is needed to ensure strong association. Furthermore, this is consistent with the dynamic, rapidly transitioning bound and unbound states of NUP153 found in our MD simulations (Figure 2E). These fast *on/off* kinetics are potentially important for the translocation of the capsid within NPC, as suggested by CA tube threading through the NuPODs (Figure 3D).

### NUP153 stabilizes capsid assemblies

How and when the capsid uncoats during HIV-1 infection remains a longstanding question (Hofer, 2020). To better understand the potential involvement of NUP153 in capsid uncoating, we tested its effect on wild-type (wt) CA tubes under different salt conditions. The wt CA tube was stable in a high salt environment such as 1M NaCl, but its lattice structure rapidly collapsed and disintegrated when the salt concentration was lowered to 75 mM (Figure 5B). However, tube disassembly at low salt concentration was averted in the presence of NUP153^1411-1475^, resulting in a large number of intact tubes coated with fuzzy irregularly-shaped NUP153^1411-1475^ proteins (Figure 5B). Similar results were obtained when testing the effect of NUP153^1411-1475^ on CA spheres using size-exclusion chromatography assays. NUP153 stabilized CA spheres under low salt conditions, preventing their disassembly (Figure S5). The stability effect of NUP153^1411-1475^ was concentration dependent. A large number of micron-long CA tubes could be observed at 50 μM NUP153, some tubes of nanometers in length remained at 5 μM NUP153, and a complete loss of tube structures resulted at 1 μM NUP153 (Figure S5). Given its high local concentration in the NPC, NUP153 binding to the capsid likely does not trigger uncoating but rather helps stabilize the lattice.

**Figure 5.**
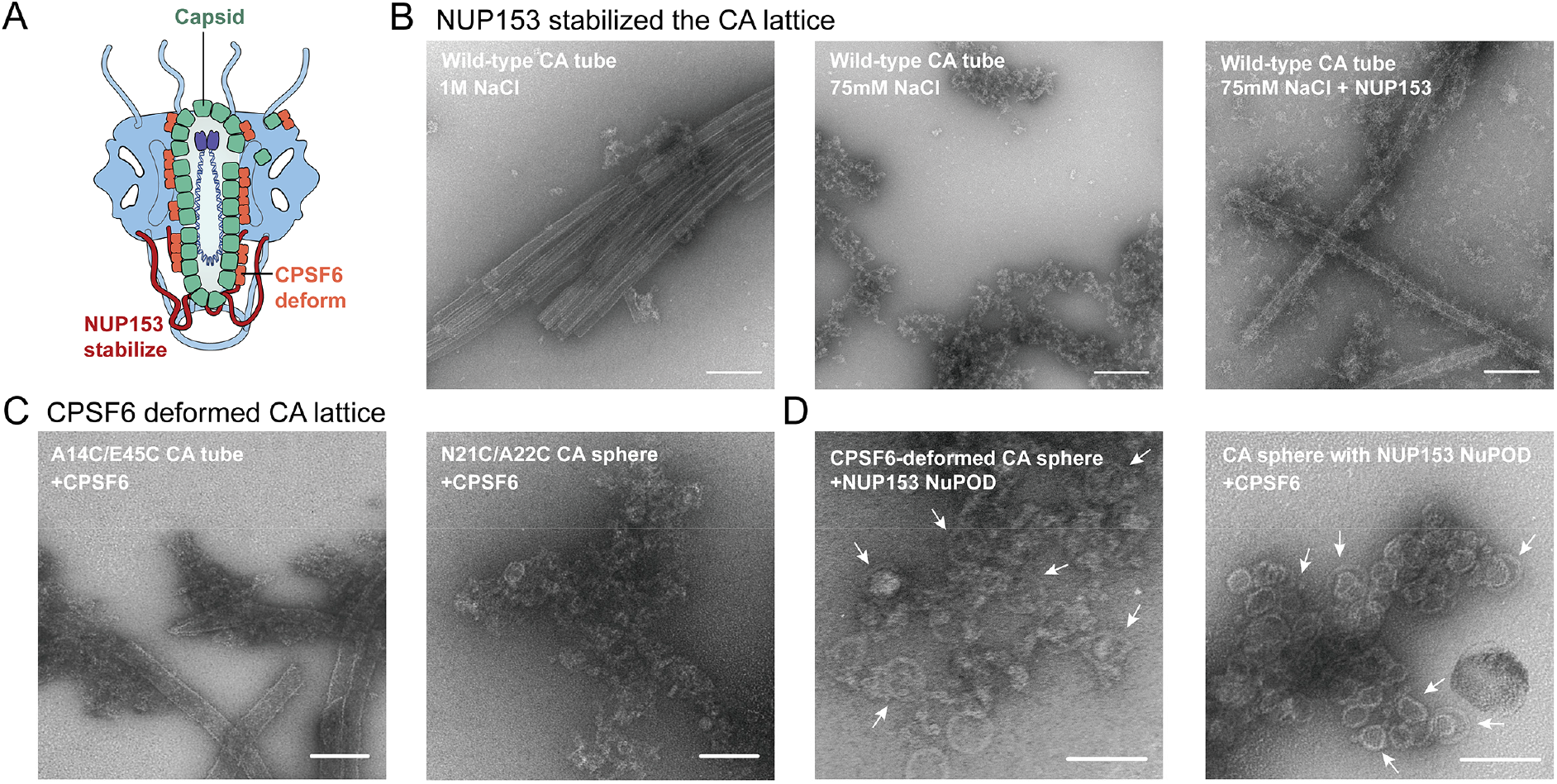
NUP153 stabilizes and CPSF6 destabilizes CA lattice. (A) Schematic of the combined effect of NUP153 and CPSF6 on the capsid. (B) Negative-stain electron micrographs of intact wt CA tubes under high salt conditions (1 M NaCl, left), disintegrated wt CA tubes under low salt conditions (75 mM NaCl, middle), and wt CA tubes stabilized by NUP153 under low salt conditions (75 mM NaCl, right). Scale bars: 200 nm. (C) CFSF6 induces crosslinked CA tubes and spheres to disassemble and/or deform. Scale bars: 100 nm. (D) Left: NUP153 NuPOD attached to CPSF6-deformed CA spheres. Right: CFSF6 was added to a pre-incubated mixture of NUP153 NuPOD with CA spheres, which remained largely intact. Scale bars: 100 nm. See also Figure S5.

### CPSF6 destabilizes capsid assemblies

One of the most intriguing aspects of HIV-1 nuclear import is the apparent dimensional mismatch between the ∼60 nm wide capsid and the ∼40 nm diameter of the nuclear pore, although heterogeneity in NPC central pore sizes has been suggested (Beck and Baumeister, 2016; Feldherr et al., 2001; Mahamid et al., 2016). This problem can potentially be resolved by the involvement of other host factors that can help remodel the capsid structure. CPSF6 facilitates capsid nuclear import and may play such a role in this process. It has been reported that truncated CPSF6 deforms and disrupts CA tubes (Ning et al., 2018). Indeed, we found that full-length CPSF6 (1 μM) efficiently disrupted CA spheres and tubes into potentially deformed CA lattices (Figure 5C and S5). NUP153 NuPODs could still bind these largely deformed spheres (Figure 5D, left), which is reminiscent of our observation of NuPOD binding to deformed native virus cores (Figure 3C, bottom right). When CA spheres were pre-incubated with the stabilizing NUP153 NuPODs, the destructive effect of CPSF6 was significantly reduced (Figure 5D, right). It is possible that the combined effect of these two host factors allows the capsid to deform yet remain largely intact to pass through NPCs.

## Discussion

HIV-1 nuclear import is one of the least understood processes of the viral life cycle. It was initially thought that the capsid immediately uncoated upon virus entry into the cell, with only recent studies pointing to the possibility of a largely intact capsid passing through the NPC into the nucleus (Burdick et al., 2020). However, mechanistic data regarding this process remain scarce. In this study we provide in-depth mechanistic insights into how HIV-1 capsid interacts with a critical NPC component, NUP153, revealing the likely import mechanism. We show that besides the known FG-motif that binds weakly to CA hexamers, NUP153 contains a previously unexplored ‘RRR’ motif at its C-terminus that is predominantly responsible for its binding to the HIV-1 capsid. Together, these two regions enable NUP153 to strongly interact with the capsid (Figure 2C). We also created a minimal 25 aa peptide composed of the FG and RRR-motifs that retained potent capsid binding *in vitro* and in the context of HIV-1 infection.

The NUP153 C-terminal region senses a pattern in capsid formed via the tri-hexamer center that only exists in the assembled capsid. This binding characteristic has profound implications on the mode by which the capsid engages the NPC to enter the nucleus. Although NUP153 was found to be able to extend to the cytosol (Fahrenkrog et al., 2002), it is part of the NPC nuclear basket likely positioned deep into the nucleus. The higher-order CA lattice targeting mode of NUP153 supports nuclear penetration of the capsid that is at least partially assembled, especially if the brunt of the many copies of NPC-associated NUP153 engage the capsid during nuclear import. These findings establish a mechanistic foundation for passage of the intact or nearly intact capsid into the nucleus (Burdick et al., 2020).

We have designed and employed NuPODs, a unique and modular experimental tool for gaining mechanistic insights into nuclear transport mechanisms. Our NuPOD system allowed us to interrogate previously unobtainable mechanistic details of HIV-1 capsid nuclear entry. For example, the NUP153 NuPOD showed an unambiguous preference to bind the tip regions of the capsid. This high-curvature preference of NUP153 potentially facilitates the correct alignment of the end of the capsid to the NPC for penetration. Rather strikingly, we captured CA tubes threading through NUP153 NuPOD rings, which may represent the scenario of capsid passing through the NPC when the central pore is of sufficient size (Figure 6). Although it is thought that the diameter of the NPC channel is less than the width of an HIV-1 capsid, recent evidence has shown that the NPC possesses a degree of elasticity, and the central channel may expand under certain conditions (Mahamid et al., 2016; Zhang et al., 2020). Consistently, our data provide direct evidence that assembled capsid structures can pass through a nuclear pore-like channel (Figure 3D).

**Figure 6.**
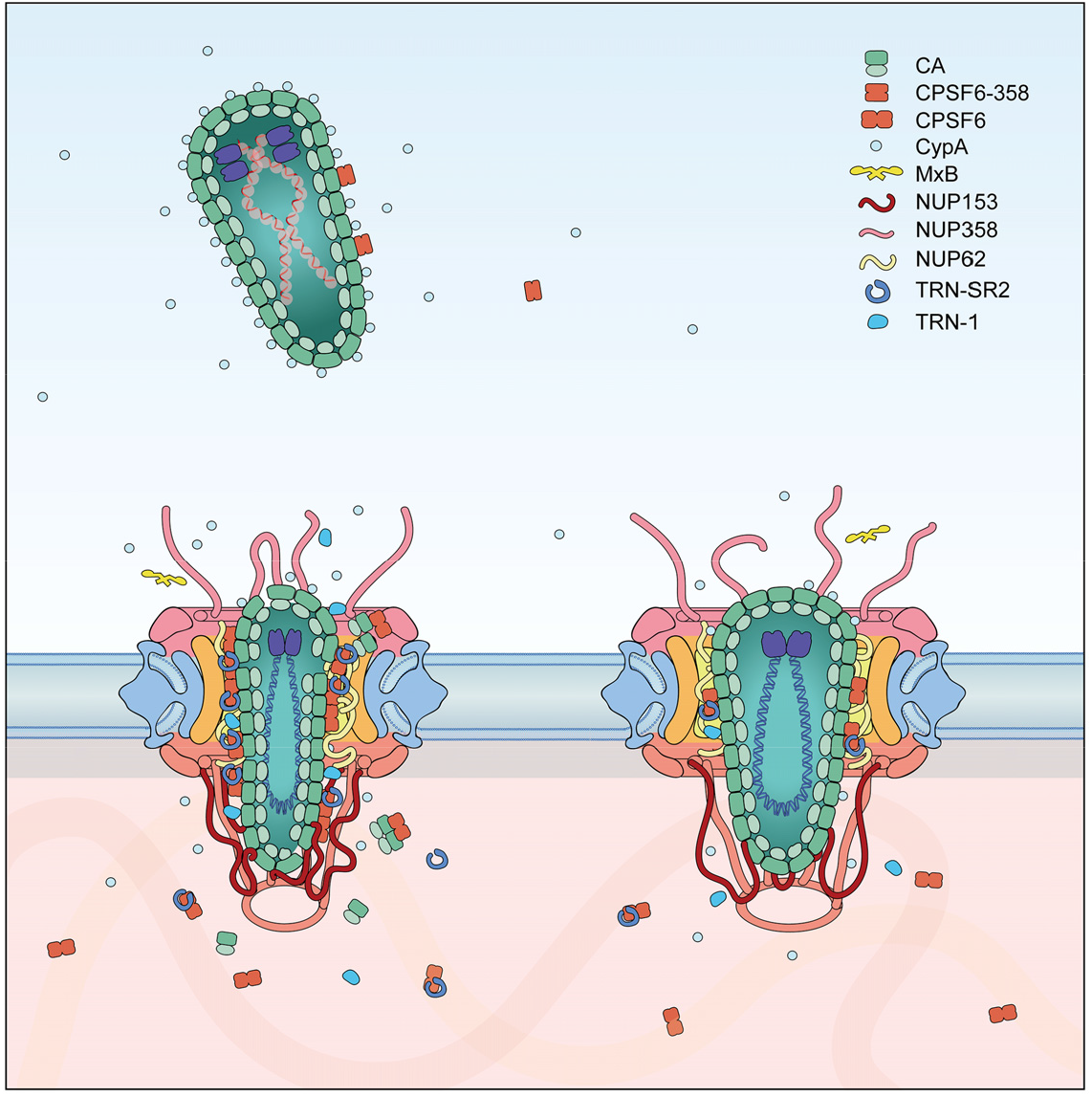
Model of the capsid passing through small (left) and large (right) NPC.

In addition to NUPs, other proteins, such as CPSF6, interact with the HIV-1 capsid to facilitate its nuclear import (Bejarano et al., 2019; Chin et al., 2015). These proteins may modulate capsid stability in opposite directions to achieve a fine balance to allow its passage through the NPC. The CPSF6-capsid interaction has also been implicated in post-entry trafficking of capsids to the interior region of the nucleus for integration (Achuthan et al., 2018; Burdick et al., 2020; Francis et al., 2020). CPSF6 deforms and causes the disintegration of capsid assemblies, as shown in our experiments and reported previously for a CPSF6 truncation variant (Ning et al., 2018). We found that NUP153 can counteract the destabilization effect of CPSF6 (Figure 5B, D). The combined effect of these two factors may enable the capsid to deform and morph through NPCs of comparatively small size and still enter into the nucleus largely intact (Figure 5D and 6). Recent studies have indicated that the capsid could remain at the nuclear pore for a few hours during its transit through the NPC (Burdick et al., 2020; Zuliani-Alvarez and Towers, 2019). One can picture a scenario whereby multiple copies of NUP153 in the NPC bind the narrow tip of the capsid to keep it positioned while CPSF6 exerts local deformations to enable the capsid structure to pass through the pore (Figure 6).

In the context of viral infection, after capsid entry into the host cell, the outcome of the infection depends on a complex network of interactions between host and viral proteins. For example, cytoplasmic CypA may attach to the capsid to protect the virus against the host immune response (Kim et al., 2019; Rasaiyaah et al., 2013), and it may also compete with NUP358 and affect capsid docking to the NPC. MxB in the vicinity of the nucleus may bind to the capsid lattice and restrict its entry into the NPC. Within the NPC and inside the nucleus, cooperation and competition between CPSF6 and other proteins such as CypA (Schaller et al., 2011) and NUP153 (Di Nunzio et al., 2012; Koh et al., 2013) may occur to enable the delivery of the capsid to sites of integration. Considering the versatility of these proteins and their interactions with each other, the network of proteins involved in the nuclear transport process is complex and intertwined (Figure 6). Perturbations in one or a subset of these proteins may lead to convoluted effects on the progress of the capsid through the NPC. This complexity presents a major hurdle for mechanistic studies of nuclear transport in cells. The NuPOD platform is a unique toolkit to enable a biochemical approach to one of the most complex aspects of cell biology. Herein, we leveraged a simple NuPOD system to gain mechanistic insight into the nuclear passage of HIV-1. With modularity and programmability, next-generation NuPODs can be built with increasing complexity to more closely mimic the environment of NPC. We envision NuPODs will continue to be an invaluable tool to study HIV-1 nuclear import as well as other viruses that rely on their capsids to gain nuclear entry (Fay and Pante, 2015; Gallucci and Kann, 2017).

## Methods

### EXPERIMENTAL MODEL AND SUBJECT DETAILS

Molecular cloning, as well as expression of recombinant proteins, was carried out in competent *E. coli* strain BL21 (DE3). Cells were cultured at 37 °C, in either Luria Broth (LB; for starter culture) or Terrific Broth (TB; for protein expression), while shaking at 220 RPM. HEK293T were cultured in Dulbecco’s modified Eagle medium supplemented to contain 10% fetal bovine serum (FBS), 100 IU/ml penicillin, and 0.1 mg/ml streptomycin at 37°C with 5% CO^2^ in a humidified incubator. HIV-1 particles were harvested from MT-4 cells infected with HIV-1 molecular clone R9. MT-4 cells were cultured in RPMI1640 medium supplemented with fetal bovine serum (10% vol/vol), penicillin (100 units/ml), and streptomycin (0.1 mg/ml) in a humidified incubator at 37°C and 5% CO^2^.

### METHOD DETAILS

#### Cloning and expression

NUP153CTD from *homo sapiens* (amino acids 896-1475), as well as yeast nucleoporin NSP1 (amino acids 2-603) from *S. cerevisiae*, was cloned as 10× His-MBP-SUMO-NUP153 (or NSP1)-SNAP constructs (Figure S1) into a pET-28a-derived vector (Novagen) and expressed in BL21-Gold (DE3). Cells were grown in TB at 37 °C with shaking until they reached an OD_600_ of 0.8. Protein expression was then induced with 1 mM IPTG for 3 hrs at 20 °C before collection by centrifugation. Cell pellets were refrigerated at −80 °C until use.

All CA constructs were cloned into pET-11a (EMD Millipore). CA-Foldon was generated by directly fusing Foldon to the C-terminus of CA. CA proteins were overexpressed in BL21-Gold (DE3) cells at 25 °C for 12 hr by induction with 0.5 mM IPTG at 0.8 OD_600_. Cell pellets were collected by centrifugation and refrigerated at −80 °C until use. Human CPSF6 was cloned into bacterial expression vector and expressed.

#### Protein purification

NUP153/NSP1-expressing cell pellets were resuspended in lysis buffer (1× PBS, 0.5 mM TCEP, 0.1 mM PMSF, 1× Roche cOmplete protease inhibitors, pH 7.4), and lysed in a homogenizer. Whole-cell lysates were then spun at 35,000 rpm for 45 min in a Type 45 Ti rotor (Beckman Coulter). Subsequently, the supernatant was decanted and filtered through a 0.45 μm cellulose acetate membrane. The resulting filtered lysate was spiked with 25 mM imidazole, then applied to a 5 mL HisTrap column (GE) on an ÄKTA system (GE) at a flow rate of 1 mL/min. The column was rinsed with a wash buffer (1× PBS, 25 mM imidazole) and eluted on a buffer gradient (1×PBS, 500 mM imidazole). Concentrations of purified protein were determined by Nanodrop (Thermo Fisher). Samples were flash-frozen in liquid nitrogen and refrigerated at −80 °C until use.

Untagged CA proteins were purified by 25 % w/v ammonium sulfate precipitation, dialysis into a low-salt buffer (25 mM HEPES, 0.1 mM TCEP, pH 7), and cation exchange chromatography. CA-foldon fusions were purified by 35 % w/v ammonium sulfate precipitation and anion exchange chromatography. All CA constructs were dialyzed into CA storage buffer before freezing or further experiments (50 mM Tris, 75 mM NaCl, 40 mM BME, pH 8). All purification steps were validated using SDS-PAGE.

#### CA tube co-pelleting assays

A14C/E45C disulfide crosslinked CA tubes were dialyzed into 25 mM Tris, pH 8 buffer. NUPs were incubated in 21 uL reactions with CA tubes for 30 min at room temperature, then spun at 20000 g for 10 min at 4 °C. Total, soluble, and pellet fractions were collected and analyzed using SDS-PAGE. NUP153CTD and all mutant constructs were incubated at 3 μM with 100 μM CA in a 25 mM Tris, 75/300 mM NaCl, pH 8 buffer.

#### Size-exclusion chromatography co-elution assays

Assembled CA hexamer, hexamer-2, and pentamer were mixed with NUP153^1411-1475^ for 30 min to 1 hr on ice in the same buffer used for CA tube co-pelleting assays (unless specified). Binding reactions were carried out in a volume of 100 uL and contained 12 μM NUPs and 72 uM monomeric concentration of CA. All binding tests were performed on GE Superdex 200 10/300 GL column in a 25 mM Tris, 75 mM NaCl, pH 8 buffer with a flow rate of 0.5 ml/min. The 280 nm absorbance was recorded.

Size-exclusion chromatography for assembled N21C/A22C CA sphere, NUP153^1411-1475^, and mixtures were performed on GE Superose 6 10/300 GL column in 25 mM Tris buffer pH 8, 50/1000 mM NaCl. The 280 nm absorbance was recorded.

#### Isothermal titration calorimetry

Binding reactions were performed in a TA Instruments NanoITC machine at 25 °C in a 25 mM Tris, 75 mM NaCl, pH 8 buffer. Hexamer-2 was stable in these conditions after overnight dialysis and during experiments. Synthesized NUP153 peptide (Genscript Co.) was resuspended in the same buffer and injected as the titrant to the Hexamer-2 in the cell. Data were analyzed using the NanoAnalyze (TA Instruments) software, and curves were fitted with an independent one-site binding model.

#### Size exclusion chromatography linked to multi-angle light scattering (SEC-MALS)

Multiangle laser light-scattering experiments were performed at room temperature in a 50 mM Tris-HCl, 150 mM NaCl, pH 8 buffer. Light-scattering data were measured using a Dawn Heleos-II spectrometer (Wyatt Technology) coupled to an Opti-lab T-rEX(Wyatt Technologies) interferometric refractometer. Samples (100 uL) at 1 mg/mL were injected and run over a Superdex 200 GL (GE Healthcare) column at a 0.5 ml/min flow rate. Data on light scattering (690 nm laser), 280 nm absorbance, and refractive index were measured. Before sample runs, the system was calibrated and normalized using the isotropic protein standard, monomeric bovine serum albumin. The dn/dc value (changes in solution refractive index with respect to protein concentration) is relatively constant for proteins (Wen, 1996), and set to 0.185 for all experiments and analysis. Data were processed in ASTRA as previously described (Wyatt, 1993).

#### Trim-NUP153 mediated restriction assays

##### Plasmid construction

TRIM-NUP153 proteins were expressed from the bicistronic expression vector pIRES2-eGFP (Shun et al., 2007). To make Trim-HA-NUP153^896-1475^, Trim and NUP153 sequences were PCR-amplified from pLPCX-Trim-HA-NUP153c (Matreyek et al., 2013) using primer pairs 5’-GATCTCGAGCTCAAGCTTCGAATTC/5’-GTAATCTGGAACATCGTATGGGTAGCCACCTCCAGATCCCCAGTAGCGTCGG3’ and 5’-CATACGATGTTCCAGATTACGCTGGAGGTGGATCTCCGCGGAACTCAGCAGCCTCCTC/5’-GATCCCGGGCCTTTATTTCCTGCGTCTAACAG, respectively (*Sac II* restriction site underlined). The linked TRIM-HA-NUP153 insert, amplified from these DNAs using primers 5’-GATCTCGAGCTCAAGCTTCGAATTC and 5’-GATCCCGGGCCTTTATTTCCTGCGTCTAACAG, was digested with *Xho I* and *Xma I* and ligated with *Xho I/Xma I*-digested pIRES2-eGFP. Trim-HA-NUP153^1401-1475^ vector was created by amplifying NUP153^1401-1475^ sequences using primers 5’-CAGCCGCGGACTACAAATTTCAACTTCACAAACAACAG and 5’-GATCCCGGGCCTTTATTTCCTGCGTCTAACAG, followed by digestion with Sac II and Xma I and swapping this DNA for the analogous region within Trim-HA-NUP153 ^896-1475^ via *Sac II* and *Xma I* restriction sites. To create Trim-HA-NUP153^1411-1475^ lacking residues of 1425-1464, double-stranded DNA representing NUP153 residues 1411-1424 and 1465-1475 was synthesized by IDT (5’CAGCCGCGGCCATCAGGAGTGTTCACATTTGGTGCAAATTCTAGCACACCTCGCAAGATAAAGACTGCTGTTAGACGCAGGAAATAAAGGCCCGGGATC3’), digested with *Sac II* and *Xma I*, and swapped for the NUP153^896-1475^ region as above. TRIM-NUP coding regions of all plasmids were verified by DNA sequencing.

##### Cells, virus production, and infection

HEK293T cells were cultured in Dulbecco’s modified Eagle medium supplemented to contain 10% fetal bovine serum (FBS), 100 IU/ml penicillin, and 0.1 mg/ml streptomycin.

Single-round HIV-1 carrying the gene for firefly luciferase (HIV-Luc), produced by transfecting HEK293T cells in 10 cm dishes with 7.5 μg pNLX.Luc.R-.ΔAvrII and 1.5 μg pCG-VSV-G using PolyJet (SignaGen Laboratories), was concentrated by ultracentrifugation as described (Serrao et al., 2016). HIV-Luc yield was assessed by p24 ELISA (Advanced Bioscience Laboratories). Cells transfected with pIRES2-eGFP expression constructs were used in HIV infection assays as previously described (Jang et al., 2019). Briefly, HEK293T cells in 6-well plates were transfected with 2 μg pIRES2-eGFP or Trim-HA-NUP153 derivatives using Effectene (Qiagen). At 24 h post-transfection, GFP-positive cells were selected in basic sorting buffer (1 mM EDTA, 25 mM HEPES, pH 8.0, 1% FBS, Mg2+/Ca2+-free phosphate-buffered saline) by fluorescence-activated cell sorting. Approximately 3-5 × 10^5^ sorted cells were lysed for immunoblotting while ∼2 × 10^5^ cells were infected in duplicate with HIV-Luc (0.25 pg p24 per cell) in the presence of 4 μg/ml polybrene. At 48 h post-infection, cells were lysed and luciferase activity was determined as previously described (Lu et al., 2004). Luciferase values were normalized to the level of total protein in cell lysates as determined by the BCA assay (Pierce).

##### Western Blotting

Details of the procedure were as described (Jang et al., 2019). 10 μg of total cellular proteins fractionated through 3-8% polyacrylamide Tris-acetate or 4-12% polyacrylamide Bis-Tris gels (Invitrogen) were subsequently transferred to polyvinylidene difluoride (PVDF) membranes. Fusion protein expression was detected with anti-hemagglutinin (HA) antibody sc7392 (Santa Cruz) followed by HRP-mouse IgG antibody (Dako); actin was detected by HRP-conjugated β-actin antibody (Abcam). Membranes developed using ECL prime reagent (Amersham Biosciences) were imaged with a ChemiDoc MP imager (Bio-Rad).

#### Molecular dynamics (MD) simulation

##### Atomic model for MD simulation

The 3-dimension structure of the last 75 residues of NUP153^1401-1475^ was built in Modeller (Eswar et al., 2006). A total number of 5,000 structures (Figure S3A) were generated and scored by the discreet optimized protein energy (DOPE) method (Shen and Sali, 2006). The candidate model of NUP153^1451-1475^ was from the structure of the NUP153^1401-1475^ with the lowest DOPE score and then subjected to a 100-ns long equilibration in explicit water in NAMD2.13 (Phillips et al., 2005). The equilibrated NUP153^1451-1475^ peptide was initially placed 1.5 nm above the tri-hexamer interface in the HIV-1 CA hexamer-2 model, which was built based on the structure of the full length native HIV-1 CA monomer (PDB:4XFX). The combined structure (Figure S3) was then solvated with TIP3P water (Jorgensen and Jenson, 1998) and neutralized by NaCl at 150 mM concentration. The resulting model had a total number of 168,500 atoms (Figure S3B).

##### MD simulation set-up

After model building, the NUP153^1451-1475^-CA hexamer-2 complex was subjected to a minimization and thermalization step. Subsequently, the whole system was equilibrated for over 100 ns at 310 K. During the equilibration, the Cα atoms of the peripheral helices of CA monomer and the N-terminal residue in NUP153^1451-1475^ were restrained. The minimization, thermalization and equilibration steps were completed in NAMD2.13 (Phillips et al., 2005). This equilibrated model then ran for a total simulation time of 8.7 μs on a special purpose computer Anton2 (Shaw et al., 2014) in the Pittsburgh supercomputing center. To mimic the CA hexamer lattice environment in the simulation, the Cα atoms of three sets of CA α-helices on the peripheries of the tri-hexamer model were applied with a harmonic restraint of 1 Kcal/mol Å^2^ in x, y and z directions (Figure S3B). Also, the Cα atoms of NUP153 residue 1451 was applied with a harmonic restraint of 1 Kcal/mol Å^2^ in z direction. CHARMM 36m (Huang et al., 2017) force field was used for all MD simulations. During the simulation, the temperature (310 K) and pressure (1 atm) was maintained by employing the Multigrator integrator (Lippert et al., 2013) and the simulation time step was set to 2.5 fs/step, with short-range forces evaluated at every time step, and long-range electrostatics evaluated at every second time step. Short-range non-bonded interactions were cut off at 17 Å; long range electrostatics were calculated using the k-Gaussian Split Ewald method (Shan et al., 2005).

##### Markov state model analysis and Model validation

Markov state models (MSMs) have been widely applied to study the conformational dynamics of biomolecules (Chodera and Noé, 2014; Husic and Pande, 2018b; Wang et al., 2018a). To identify the important metastable states of NUP153CTD in the MD simulation, Markov state models were constructed and validated in the PyEMMA 2.5.7 package (Scherer et al., 2015). First, the coordinates of NUP153^1451-1475^ from the Anton2 simulation were directly transformed into two features, the Cartesian coordinates of NUP153^1451-1475^ backbone atoms and 4,853 distance pairs between the backbone atoms of NUP153^1451-1475^. Subsequently, time-lagged independent component analysis (TICA) (Pérez-Hernández et al., 2013) was performed to decompose these features onto 100 slow independent components (ICs) (Figure S3D). The projected data set was then discretized using the k-means method (MacQueen, 1967) and resulted in fifty microstates. The transition probability matrix between these microstates was then computed at a selected lag time of 1.2 ns (Figure S3J). Finally, an eight-state MSM using the PCCA+ algorithm (Röblitz and Weber, 2013) was constructed and the mean first passage times (MFPTs) between different states were estimated (Figure S3I). The eight-state MSMs built were validated with the Chapman-Kolmogorov test (Noe et al., 2009; Prinz et al., 2011).

#### DNA origami assembly

DNA origami structures were designed in caDNAno (caDNAno.org). ssDNA handles were extended from the 3’ end of staple strands at positions indicated in Figure 3B. The handle sequences were 5’-AAATTATCTACCACAACTCAC-3’ for inner/outer handles. These structures were assembled using an M13mp18 bacteriophage-derived circular ssDNA strand (8064 nt) and oligonucleotides from Integrated DNA Technologies over 36 hr on an 85–25 °C annealing gradient. Structures were purified using rate-zonal centrifugation through a 15–45 % glycerol gradient in a 1× TE, (5 mM Tris-Cl, 1 mM EDTA), 16 mM MgCl_2_, pH 8.0 buffer in an SW 55 rotor (Beckman Coulter) at 50,000 rpm at 4°C for 1 hr, followed by fraction collection (Lin et al., 2013). Fractions containing purified DNA origami structures were buffer exchanged into a 1× TE, 16 mM MgCl_2_, pH 8.0 buffer and stored at −20 °C.

#### Benzylguanine (BG)-DNA preparation

5’-labeled amino-DNA oligonucleotides (anti-handles; IDT) were resuspended in de-ionized H_2_O at 2 mM concentration. BG-NHS was dissolved in DMSO at 20 mM. Anti-handles and BG-NHS were mixed in a 1:3 ratio in 67 mM HEPES, pH 8.5 buffer, and incubated at room temperature for 1 hour. BG-DNA was then purified from excess BG-NHS by ethanol precipitation. Dried BG-DNA pellets were stored at −20 °C until use.

#### Protein-DNA conjugation

BG-DNA pellets were resuspended in de-ionized H_2_O and mixed with purified NUP153CTD in 1× PBS buffer at a final concentration of 40 μM BG-DNA and 20 μM NUP153CTD (2:1). This reaction was incubated at 25 °C for 2 hr. Excess DNA was removed from conjugated proteins using size exclusion chromatography on a Superdex 200 10/300 GL column (GE Healthcare) in a 25 mM Tris, 75 mM NaCl, pH 8 buffer. Conjugation efficiency was verified by SDS-PAGE using Coomassie or SYPRO Red stain (Thermo Fisher).

#### Hybridization of NUP to DNA origami

DNA-conjugated NUP153CTD was added to DNA origami rings at 1.5× excess over the number of origami handles (e.g. 5 nM origami × 32 handles × 1.5 = 240 nM FG-NUP153-DNA) in a 1× TE, 16 mM MgCl_2_, pH 8.0 buffer and incubated for 2 hr at 37 °C. NUP153 NuPODs were purified by rate-zonal centrifugation, as described in “DNA Origami Assembly”, through a 15–45 % glycerol gradient in the hybridization buffer. The purified product is called a NUP153 NuPOD (NucleoPorins Organized on DNA).

#### CA assembly

##### Hexamer-2, hexamer and pentamer

Hexamer-2 was assembled using a 1:1 molar ratio of the appropriate CA proteins (Summers et al., 2019), resulting in 10-40 mg/mL of total proteins. Mixtures were dialyzed overnight (using Thermo Slide-a-lyzer dialysis cassettes) in a 50 mM Tris, 1 M NaCl, pH 8 buffer. Mixtures were dialyzed for a second night in a 50 mM Tris, pH 8 buffer. Hexamer-2 assemblies eluted from an anion exchange column at approximately 250 mM NaCl. Each assembly was purified using a Superdex 200 PG or Superdex 200 GL size-exclusion chromatography column (GE Healthcare) ran with a 50 mM Tris, 300 mM NaCl, pH 8 buffer. Hexamer (A14C/E45C/W184A/M185A) and pentamer (N21C/A22C/W184A/M185A) assemblies were dialyzed (using Thermo Slide-a-lyzer dialysis cassettes) for 48 hr in a 50 mM Tris, pH 8 buffer without redox reagents. Then the crosslinked hexamers or pentamers were purified using a Superdex 200 GL column. All assemblies were concentrated to 20-50 mg/mL and frozen at −80 °C for storage.

##### Crosslinked CA tube

A14C/E45C disulfide crosslinked CA tubes were assembled by dialyzing (in Slide-A-Lyzer dialysis cassettes) CA A14C/E45C into a 50 mM Tris, 1 M NaCl, pH 8 buffer at 15 mg/mL for one night, followed by dialysis into 50 mM Tris for another night. The crosslinked tube was stored in a 50 mM Tris, 50 mM NaCl, pH 8 buffer and could remain stable for months.

##### Wild-type CA tube

Wild-type CA tubes were assembled by dialyzing into a 50 mM Tris, 1 M NaCl, pH 8 buffer at 5 mg/mL for three to four nights. Assemblies were stored in a 50 mM Tris, 1 M NaCl, pH 8 buffer.

##### CA sphere

Following a previously established protocol (Zhang et al., 2018), purified N21C/A22C CA proteins (1 mg/mL) were dialyzed into a 50 mM Tris-HCl, 1 M NaCl, pH 8.0 buffer overnight at 4°C.

##### CA cone

A14C/E45C CA protein was diluted in a 25 mM MES, 100 mM NaCl, pH 6.0 buffer to 200 μM, then mixed with a 20 to 40 fold molar excess of IP6 in solution. The assembly reaction was maintained at room temperature for 5 min until the appearance of cloudy CA assemblies. 200 uM CypA was then added to reduce the aggregation of the cones. The CA assemblies were diluted 20 times before observation under negative-stain TEM.

#### Isolation of HIV-1 cores

Native HIV-1 cores were prepared from concentrated HIV-1 particles by centrifugation through a layer of 0.5% Triton X-100 into a linear sucrose density gradient as described (Shah and Aiken, 2011). For this purpose, HIV-1 particles were harvested from 200 ml cultures of MT-4 cells (Pauwels et al., 1987) infected with undiluted virus harvested from 10-cm dishes transfected with the wild type HIV-1 molecular clone R9 (Gallay et al., 1997), which is nearly identical to NL4-3 in its protein coding sequences. To produce the virus, cells were cultured in RPMI1640 medium supplemented with fetal bovine serum (10% vol/vol), penicillin (100 units/ml), and streptomycin (0.1 mg/ml) in a humidified incubator at 37°C and 5% CO^2^. For virus production in MT-4 cells, one hundred million cells were pelleted, resuspended in 50 ml of fresh medium, and inoculated with a quantity of HIV-1 corresponding to 5000 ng of p24 in the presence of DEAE-dextran (20 mg/ml) to promote infection. Cells were subsequently cultured for 24 h, followed by removal of the inoculum by pelleting and aspiration. The cell pellet was resuspended in 200 ml of fresh medium and cultured for 4 to 5 days. The extent of infection was assessed daily by monitoring the culture visually for cytopathic effects. At approximately 5 days after inoculation, the cells were pelleted by centrifugation and the supernatants collected and clarified by passing through a 0.45 mM pore-size vacuum filter. The HIV-1 particles were concentrated by ultracentrifugation, resuspended, and HIV-1 cores were isolated. The quantity of viral CA protein in the dense gradient fractions corresponding to HIV-1 cores was determined by p24 ELISA (Wehrly and Chesebro, 1997). The presence of intact HIV-1 cores in these fractions was ascertained by negative stain transmission electron microscopy using uranyl formate. Fractions containing HIV-1 cores were aliquoted and flash frozen in liquid nitrogen and stored at −80°C until use.

#### Negative-stain electron microscopy

Capsid assemblies with origami samples were deposited onto glow-discharged 400 mesh formvar/carbon-coated copper grids (Electron Microscopy Sciences). Grids were then stained with 2% uranyl formate. Imaging was performed on a JEOL JEM-1400 Plus microscope operated at 80 kV with a bottom-mount 4k×3k CCD camera (Advanced Microscopy Technologies).

#### Statistical analysis

The data analysis was performed using the SPSS 26.0 software package (IBM, United States). All data were expressed in mean ± standard deviation (SD) if not specifically addressed. For the co-pelleting assay, differences were tested by Student’s t-test with variances verified by Fisher’s F-test. NuPOD binding counting was analyzed by the Chi-Squared goodness of fit test. P < 0.05 were considered statistically significant.

## SUPPLEMENTAL INFORMATION

## ACKNOWLEDGMENTS

We thank the Xiong Lab for discussion. This work was supported by NIH grants P50AI150481 (Y.X., C.A., and A.N.E.), P30GM110758 (J.R.P.), R01AI052014 (A.N.E.), T32AI007386 (G.J.B.), R21GM109466 (C.L. and C.P.L.), R01GM105672 (C.P.L.), R01GM132114 (C.L.) and Collaboration Development Pilot Program awards from the Pittsburgh Center for HIV Protein Interactions (J.R.P. and C.L.), and a Singapore Agency for Science, Technology and Research Graduate Scholarship (Q.X.). Anton 2 computer time was provided by the Pittsburgh Supercomputing Center (PSC) through NIH Grant R01GM116961. The Anton 2 machine at PSC was generously made available by D.E. Shaw Research.

## AUTHOR CONTRIBUTIONS

Conceptualization, Q.S., C.L., and Y.X.; Methodology, Q.S, C.X., S. J., C.A., J.R.P., A.N.E., C.P.L., C.L., Y.X.; Investigation; Q.S., C.X., S. J., Q.X., S.C.D., T.T., G.J.B., T.N.T., Y.H., S.Y., J.T., J.S., C.A., J.R.P., A.N.E., C.P.L., C.L., Y.X.; Writing–Original Draft, Q. S.; Writing – Review & Editing, Q.S, C.X., S. J., Q.X., S.C.D., J.T., C.A., J.R.P., A.N.E., C.P.L., C.L., and Y.X.; Funding Acquisition, C.A., J.R.P., G.J.B., A.N.E., C.P.L., C.L., and Y.X..

