## Supplemental information for "A DNA-origami nuclear pore mimic reveals nuclear entry mechanisms of HIV-1 capsid"

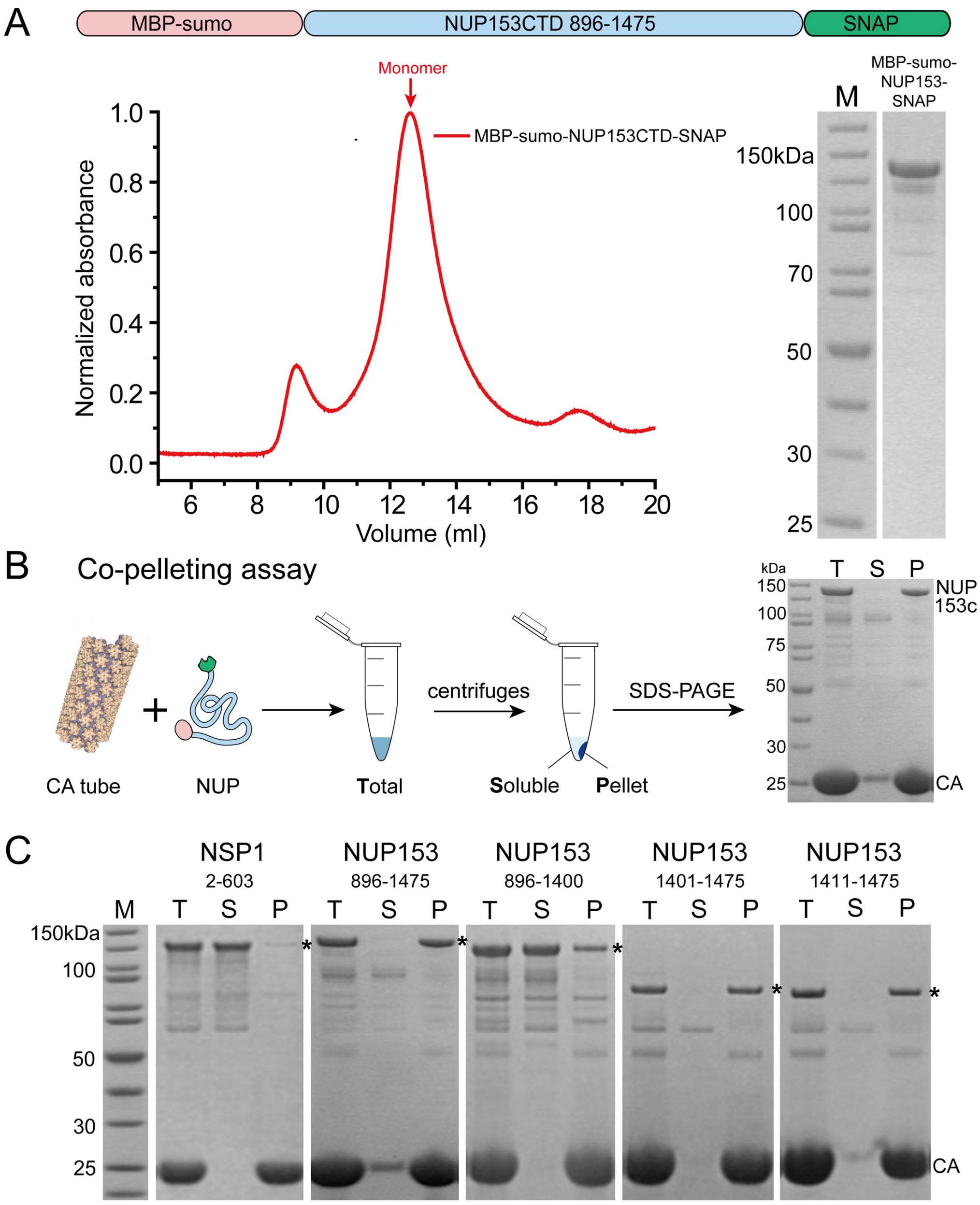


**Figure S1. NUP153CTD purification and CA tube co-pelleting assay**
(A) Top: Schematic of the MBP-sumo-NUP153-SNAP construct. Bottom-left: Size exclusion chromatography analysis shows that NUP153 is monodisperse in solution. Bottom-right: SDS-PAGE analysis of protein purity.
(B) Schematic of the CA tube co-pelleting assay.
(C) Total (T), Soluble (S), and Pellet (P) fractions of co-pelleting assays using A14C/E45C disulfide crosslinked CA tubes analyzed by SDS-PAGE; *, NUP.

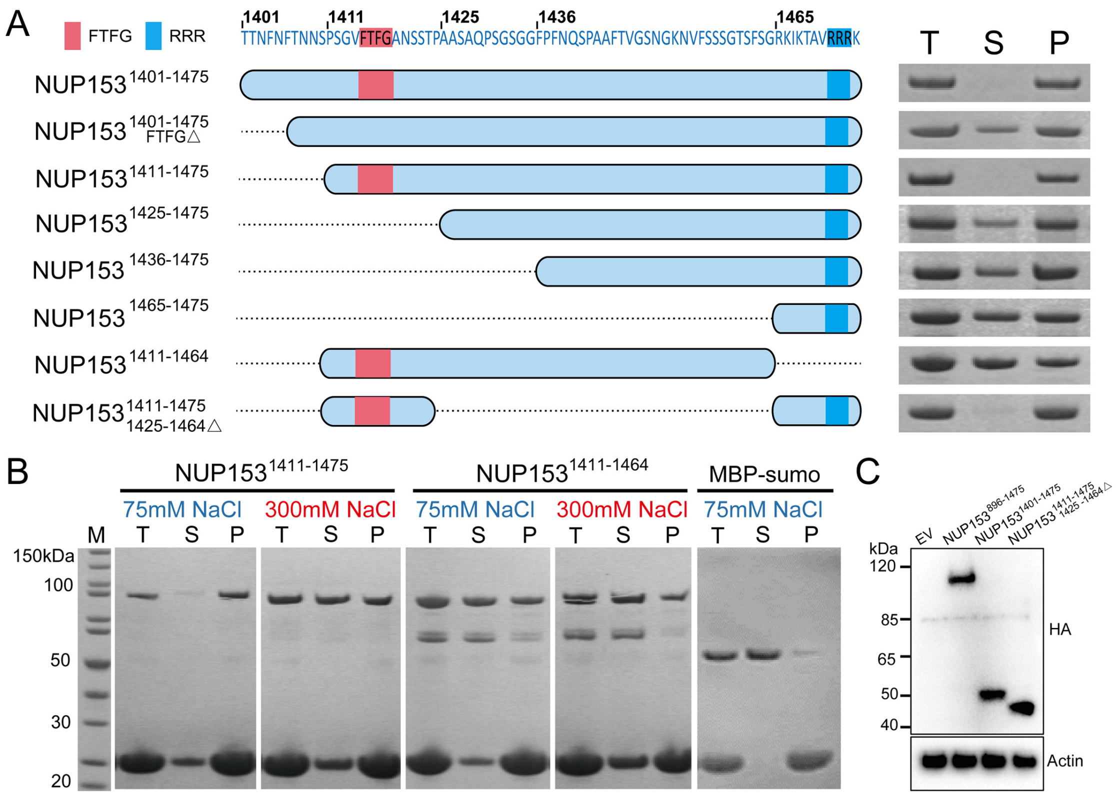


**Figure S2. Co-pelleting assay of NUP153 variants with CA tubes and protein expression in cells.**

(A) Co-pelleting assay of NUP153 variants with CA tubes. Middle: Schematics of various truncated NUP153CTD segments. Right: Total (T), Soluble (S), and Pellet (P) fractions of co-pelleting assays using A14C/E45C disulfide crosslinked CA tubes analyzed by SDS-PAGE.

(B) Results of the co-pelleting assay under low-salt and high-salt conditions.

(C) Immunoblot showing expression of NUP153CTD constructs in HEK293T cells; β-actin was blotted as a loading control.

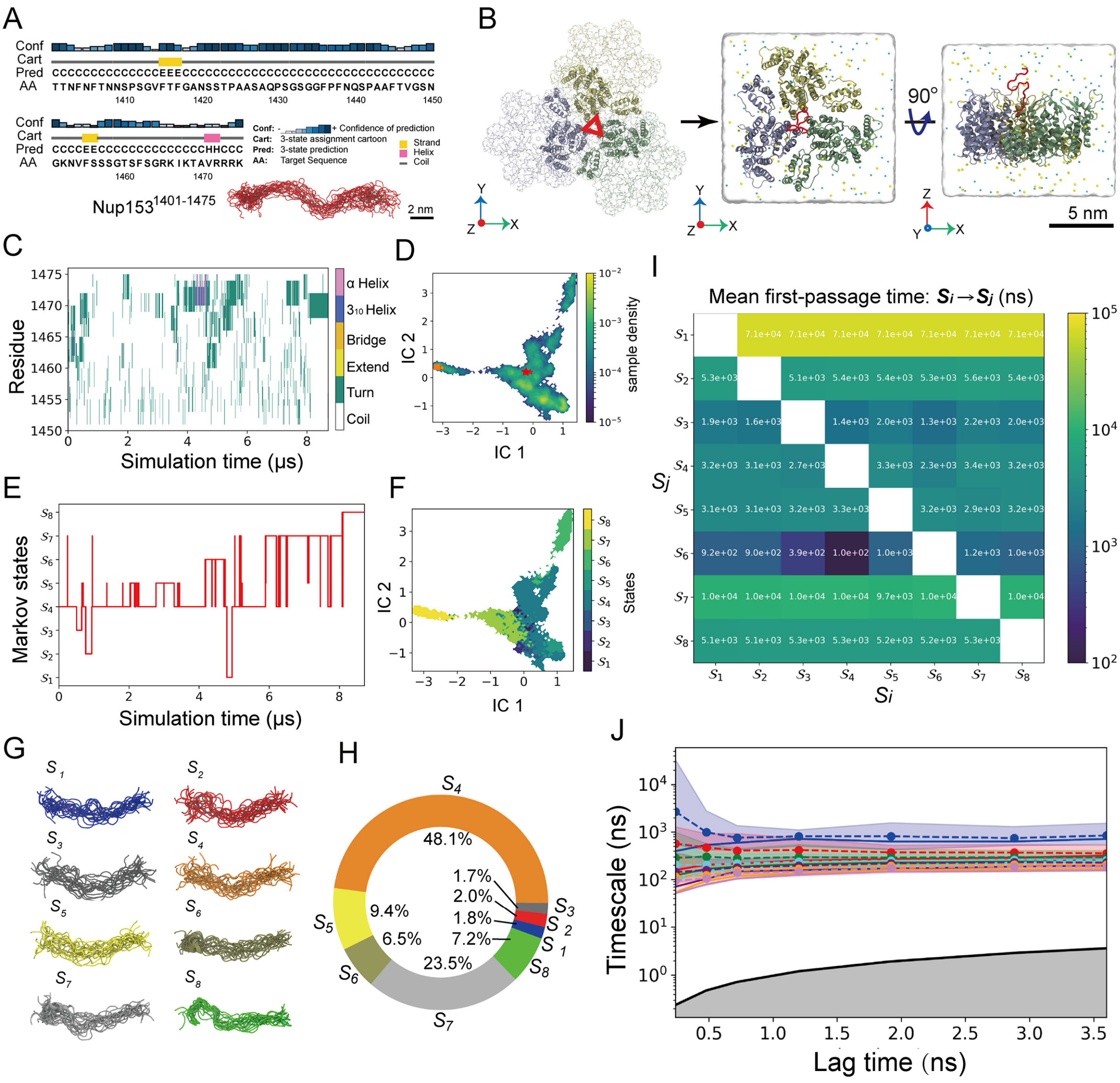


**Figure S3. The conformational dynamics of NUP153$^{1451\text{-}1475}$ at the CA tri-hexamer interface.**

(A) Modeling of the NUP153$^{1401\text{-}1475}$ structure. Left, the sequence of NUP153$^{1401\text{-}1475}$ and its PSIPRED secondary structure prediction (McGuffin et al., 2000). Right, an ensemble of the top 25 NUP153$^{1401\text{-}1475}$ models scored by the discreet optimized protein energy (DOPE).

(B) A NUP153$^{1401\text{-}1475}$-CA hexamer-2 complex. The CA dimers from different hexamers are colored in light blue, light green, and yellow, respectively, while the NUP153 peptide is shown as a red ribbon.

(C) The evolution of NUP153$^{1451\text{-}1475}$ secondary structures during the MD simulation, which were computed by the STRIDE program (Frishman and Argos, 1995) in VMD.

(D) The distribution of NUP153$^{1451\text{-}1475}$ conformations projected onto the first two independent components (ICs) space. The red star and orange square mark the starting and ending conformations of NUP153$^{1451\text{-}1475}$ in the MD simulation.

(E) Conformational transitions and (F) distributions of NUP153$^{1451\text{-}1475}$ during the simulation. Eight Markov states of NUP153$^{1451\text{-}1475}$ in the CA tri-hexamer interface were identified.

(G) Twenty representative NUP153 structures and (H) populations of the Markov states are illustrated.

(I) The mean free-passage times (MFPTs) of transitions between Markov states.

(J) Optimization of the lag-time for constructing Markov state models (MSMs). The seven slowest relaxation timescales of the MSMs as a function of the lag-time are shown. The MSM results reported in the present study were obtained with a lag-time of 1.2 ns, after which the timescales are nearly constant.

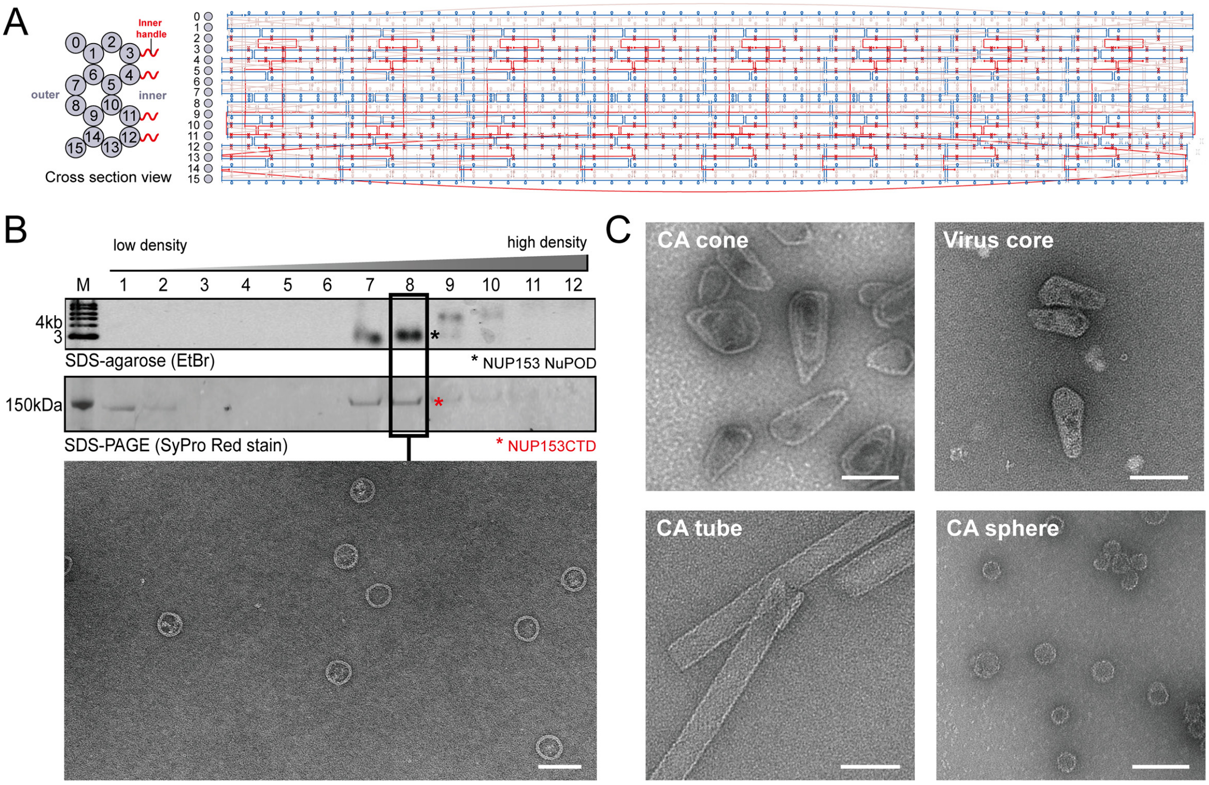


**Figure S4. NUP153 NuPOD, CA assemblies, and virus core preparation.**

(A) DNA origami designs. Cross section view (left) and strand diagrams for the 0-32 inner handle NuPOD (right). Scaffold strand is in blue; staples for inner handles are shown in red.

(B) Rate-zonal centrifugation purified NUP153 hybridized NuPOD. SDS-PAGE and SDS-agarose gel shows that NuPOD assembled with NUP153CTD and enriched in fraction 8, as confirmed by negative-stain TEM. Scale bar: 100 nm.

(C) Negative-stain electron micrographs of CA assemblies and native virus cores. Scale bar: 100 nm

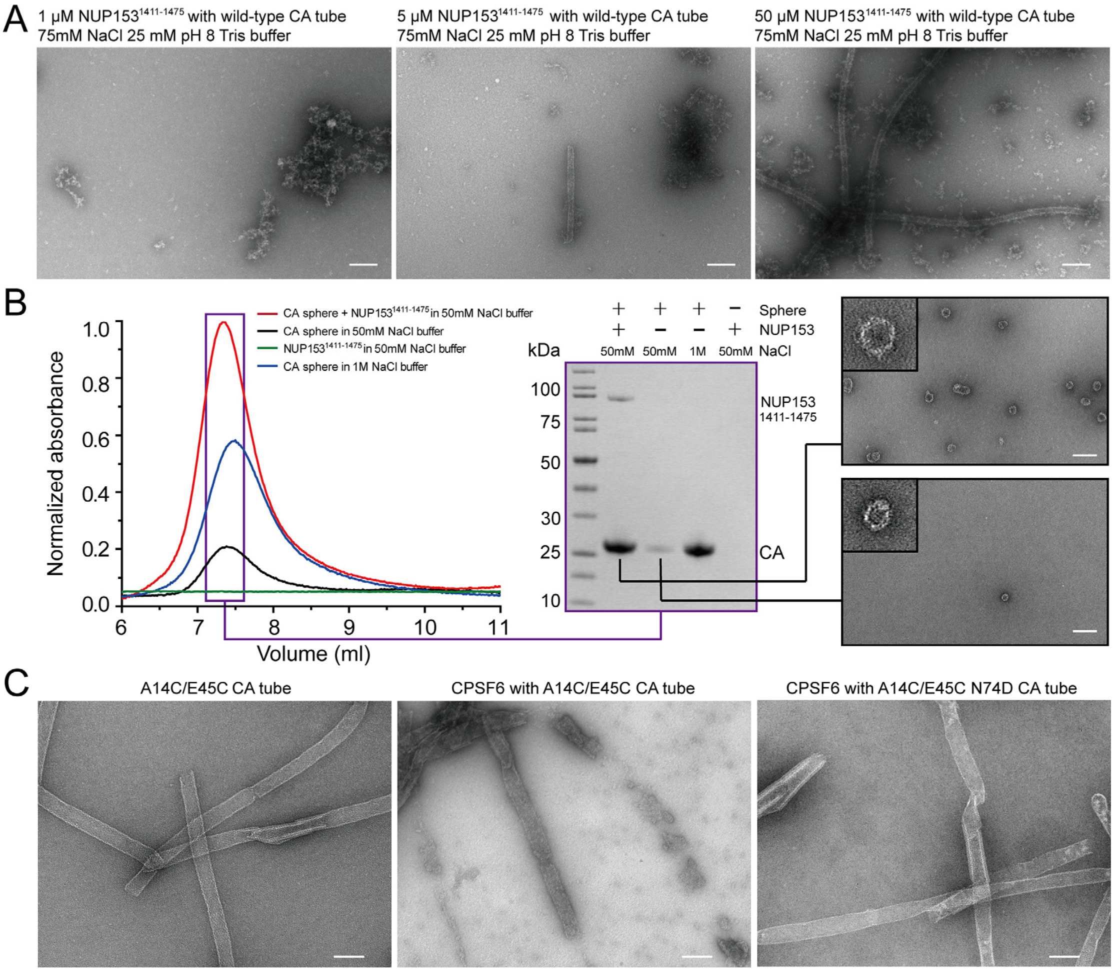


**Figure S5. NUP153 stabilizes and CPSF6 destabilizes CA lattice.**

(A) Different concentrations of NUP153$^{1411\text{-}1475}$ (1, 5, 50 μM) incubated with wild-type CA tubes. Scale bar: 200 nm.

(B) Analysis of the CA sphere under high salt (1 M NaCl) and low salt (50 mM NaCl) conditions. Left: SEC of CA spheres under high or low salt conditions with or without NUP153$^{1411\text{-}1475}$. Middle: SDS-PAGE analysis of collected SEC peak fractions. Right: negative-stain electron micrographs of CA spheres in the presence (top-right) or absence (bottom-right) of NUP153 under low salt (50 mM NaCl) conditions. Scale bars: 100 nm.

(C) A14C/E45C CA tube without CPSF6 (left); A14C/E45C CA tube with 1μM CPSF6 (middle); A14C/E45C N74D CA tube with 1 μM CPSF6 (right). Scale bar: 100 nm.

Table S1. **NUP153$^{1451-1475}$-CA hexamer-2 inter-residue contacts categorized by Markov states.**

| Markov states | NUP153 residues | CA residues | Occupancy (%) |
|---|---|---|---|
| State 1 | GLY1464 | ARG82 | 100.0 |
| | ARG1465 | ASP81 | 98.4 |
| | GLY1464 | ASP81 | 96.9 |
| | SER1463 | ASP81 | 93.8 |
| | SER1463 | ARG82 | 93.8 |
| State 2 | ARG1465 | TRP80 | 100.0 |
| | ARG1465 | LEU83 | 98.6 |
| | ARG1465 | HIS84 | 97.3 |
| | PHE1462 | PRO125 | 95.9 |
| | ILE1467 | LEU83 | 94.5 |
| | ARG1465 | GLU98 | 94.5 |
| | PHE1462 | ILE129 | 91.8 |
| | GLY1464 | TRP80 | 90.4 |
| State 3 | PHE1462 | PRO99 | 96.7 |
| | PHE1462 | PRO125 | 96.7 |
| | PHE1462 | ILE129 | 96.7 |
| | PHE1462 | GLU98 | 95.1 |
| | LYS1466 | GLU79 | 95.1 |
| | LYS1466 | GLU76 | 95.1 |
| | ARG1474 | ARG82 | 95.1 |
| | PHE1462 | ILE124 | 91.8 |
| | PHE1462 | VAL126 | 90.2 |
| | ILE1467 | LEU83 | 90.2 |
| | GLY1464 | ILE129 | 90.2 |
| | ARG1473 | ARG82 | 90.2 |
| State 6 | ILE1467 | ARG82 | 92.4 |
| | THR1469 | ARG82 | 90.7 |
| State 8 | ILE1467 | ASP81 | 100.0 |
| | ARG1472 | THR210 | 98.9 |
| | LYS1466 | ARG82 | 98.1 |
| | ILE1467 | ARG82 | 98.1 |
| | ARG1473 | GLU213 | 95.8 |
| | ARG1473 | GLY208 | 94.6 |
| | ILE1467 | ALA78 | 93.5 |
| | ALA1470 | ASN74 | 92.0 |

Only contacts with occupancies greater than 90% were shown. In Markov state 4, 5, and 7, NUP153 was found to have no residue contact with an occupancy greater than 90% with CA. The NUP153-CA residue contacts were defined as the NUP153-CA inter-residue distance smaller than a cutoff distance (3.4 Å). The contact occupancy was calculated by $100\% \times N_{ij}/n$, where $N_{ij}$ is the number of occurrences of a residue contact between NUP153 residue 'i' and CA residue 'j' in the simulation and 'n' is the total number of the MD simulation snapshots.

**Table S2. Primers used to generate NUP153 variants, related to STAR Methods**

| Primer | Sequence |
|---|---|
| NUP153-F | ATGCCATATGGAAAATTTATACTTCCAAGGTAACTCAGCAGCCTCCTCATCCTTCA |
| NUP153-R | ATGCCTCGAGTCAGCAAAGCTTTTTCCTGCGTCTAACAGCAGTCTTT |
| SNAP-F | ATGTGCCCAAGCTTATGGACAAAGATTGCGAAATGAAACGTACCACCC |
| SNAP-R | ATGCCGCTCGAGTCATCCCAGACCCGGTTTACCCAGACGA |
| NUP153 1475 F | ATGGACAAAGATTGCGAAATGAAACGTAC |
| NUP153 1400 R | GCTGCTGCCAAACTGGAAAGCCGAA |
| NUP153 1401 F | ACTACAAATTTCAACTTCACAAACAACAGTCCATCAG |
| NUP153 1411 F | CCATCAGGAGTGTTCACATTTGGTGCAAATTC |
| NUP153 1425 F | GCAGCCTCAGCCCAGCCTTC |
| NUP153 1436 F | TTTCCATTTAACCAGTCTCCAGCAGCAT |
| NUP153 1465 F | CGCAAGATAAAGACTGCTGTTAGACGCA |
| NUP153 896 R | ACCTTGGAAGTATAAATTTTCCATATGTCCACCAATCT |
| NUP153 1424 R | AGGTGTGCTAGAATTTGCACCAAATGTG |
| NUP153 1464 R | ACCAGAGAATGAAGTTCCAGAAGAAGAGA |
| NUP153 1415FTFG F | AACAGTCCATCAGGAGTGGCAAATTCTAGCACACC |
| NUP153 1415FTFG R | GGTGTGCTAGAATTTGCCACTCCTGATGGACTGTT |
| Trim F | GATCTCGAGCTCAAGCTTCGAATTC |
| Trim R | GTAATCTGGAACATCGTATGGGTAGCCACCTCCAGATCCCCAGTAGCGTCGG |
| NUP153 F | CATACGATGTTCCAGATTACGCTGGAGGTGGATCTCCGCGGAACTCAGCAGCCTCCTC |
| NUP153 R | GATCCCGGGCCTTTATTTCCTGCGTCTAACAG |
| Trim-HA-NUP153 F | GATCTCGAGCTCAAGCTTCGAATTC |
| Trim-HA-NUP153 R | GATCCCGGGCCTTTATTTCCTGCGTCTAACAG |
| NUP153 1401-1475 F | CAGCCGCGGACTACAAATTTCAACTTCACAAACAACAG |
| NUP153 1401-1475 R | GATCCCGGGCCTTTATTTCCTGCGTCTAACAG |
| NUP153 25AA | CAGCCGCGGCCATCAGGAGTGTTCACATTTGGTGCAAATTCTAGCACACCTCGCAAGATAAAGACTGCTGTTAGACGCAGGAAATAAAGGCCCGGGATC |
